# Subcellular architecture and metabolic connection in the planktonic photosymbiosis between Collodaria (radiolarians) and their microalgae

**DOI:** 10.1101/2021.03.13.435225

**Authors:** Johan Decelle, Giulia Veronesi, Charlotte LeKieffre, Benoit Gallet, Fabien Chevalier, Hryhoriy Stryhanyuk, Sophie Marro, Stéphane Ravanel, Rémi Tucoulou, Nicole Schieber, Giovanni Finazzi, Yannick Schwab, Niculina Musat

**Affiliations:** Univ. Grenoble Alpes, CNRS, CEA, INRAe, IRIG-LPCV, Grenoble, France; Helmholtz Centre for Environmental Research – UFZ, Department of Isotope Biogeochemistry, Leipzig, Germany; CNRS, Laboratoire de Chimie et Biologie des Métaux (LCBM) UMR 5249 CNRS-CEA-UGA, F-38054 Grenoble, France and CEA, LCBM, F-38054 Grenoble, France and Université Grenoble Alpes, LCBM, F-38054 Grenoble, France; ESRF, The European Synchrotron, 71, Avenue des Martyrs, 38043 Grenoble, France; Institut de Biologie Structurale (IBS), University Grenoble Alpes, CEA, CNRS, 38044 Grenoble, France; Sorbonne Universités, UPMC Université Paris 06, CNRS, Laboratoire d’Océanographie de Villefranche (LOV) UMR7093, Observatoire Océanologique, 06230 Villefranche-sur-Mer, France; Cell Biology and Biophysics Unit, European Molecular Biology Laboratory (EMBL), 69117 Heidelberg, Germany

**Keywords:** photosymbiosis, oceanic plankton, microalgae, 3D electron microscopy, subcellular imaging, dinoflagellate, radiolarians

## Abstract

Photosymbiosis is widespread and ecologically important in the oceanic plankton but remains poorly studied. Here, we used multimodal subcellular imaging to investigate the photosymbiosis between colonial Collodaria and their microalga dinoflagellate (*Brandtodinium*) collected in surface seawaters. We showed that this symbiosis is a very dynamic system whereby symbionts interact with different host cells via extracellular vesicles within the “greenhouse-like” colony. 3D electron microscopy revealed that the volume of the photosynthetic apparatus (plastid and pyrenoid) of the microalgae increased in symbiosis compared to free-living while the mitochondria volume was similar. Stable isotope probing coupled with NanoSIMS showed that carbon and nitrogen were assimilated and stored in the symbiotic microalga in starch granules and purine crystals, respectively. Nitrogen was also allocated to the algal nucleus (nucleolus). After 3 hours, low ^13^C and ^15^N transfer was detected in the host Golgi. Metal mapping revealed that intracellular iron concentration was similar in free-living and symbiotic microalgae (ca 40 ppm) and two-fold higher in the host, whereas copper concentration increased in symbiotic microalgae (up to 6900 ppm) and was detected in the host cell and extracellular vesicles. Sulfur mapping also pinpointed the importance of this nutrient for the algal metabolism. This study, which revealed subcellular changes of the morphology and nutrient homeostasis in symbiotic microalgae, improves our understanding on the metabolism of this widespread and abundant oceanic symbiosis and paves the way for more studies to investigate the metabolites exchanged.

## Introduction

In the surface layer of oceanic waters, planktonic organisms display a wide range of trophic modes to access energy, such as intimate symbiotic partnerships between taxonomically and physiologically different cells (1). While increasing connectivity and complexity of food webs, planktonic symbioses contribute to carbon and nitrogen fixation, predation, and biogeochemical cycling of different elements (2–4). The increasing recognition of the ecological role of planktonic symbioses in the global ocean stresses the need to study their physiology and underlying mechanisms that remain largely unexplored. This knowledge gap is because the large majority of planktonic symbioses are not amenable to *ex situ* laboratory culture and their genomes and life cycle are unknown. One of the most prevalent eukaryotic associations in the open ocean is the photosymbiosis between the unicellular radiolarian hosts (Rhizaria) and their intracellular symbiotic microalgae (e.g. haptophyte or dinoflagellate) (5–7). Alike coral reefs, it is a mutualistic symbiosis whereby microalgae provide photosynthetically-derived products to the host, which in turn maintains a sheltered and relatively nutrient-rich microhabitat for their microalgae. The symbiosis therefore provides a competitive advantage in nutritionally-demanding habitats, such as oceanic waters where nutrients (e.g. nitrogen, iron) are extremely poorly available. In the association between the host Acantharia (radiolarians) and the microalga *Phaeocystis* (haptophyte), the microalga is morphologically transformed into a powerful photosynthetic machinery (e.g. multiplication of voluminous plastids and larger C-fixing pyrenoids), and its nutrient homeostasis is significantly altered (e.g. trace metals) (8,9). This partnership would correspond to algal farming where the host has a strong control over its microalgae. It is not known whether similar mechanisms and such algal transformation are common in other oceanic photosymbioses involving different hosts and microalgal symbionts. In the context of evolution, photosymbioses between single-celled hosts and intact microalgae can be considered as a transitional step in plastid acquisition in eukaryotes, representing the best available experimental systems to understand how a host cell can control and benefit from the metabolism of a photosynthetic cell before gene transfer and genetic control.

Colonial radiolarians (Collodaria) are arguably one of the most abundant photosymbioses in the oceanic plankton. Collodaria were also among the first observations of symbiotic associations in nature (i.e. T.H. Huxley during the Rattlesnake expedition in 1847). Collodaria, composed of several cells, are known to host thousands of microalgae (the dinoflagellate *Brandtodinium nutricula*) in their large gelatinous matrix (10,11). Large-scale environmental DNA sequencing projects (e.g. Tara-*Oceans*) showed that collodarians are widely distributed in the oceans, and tend to numerically dominate the plankton community in surface oligotrophic waters (7,12). In tropical and subtropical waters, their total biomass can be as important as the one of the traditional zooplankton (13). High densities of colonies (16,000 - 20,000 colonies per m^3^) have been reported in different oceanic regions, with marked seasonal episodes of blooming (Khmeleva, 1967; Caron and Swanberg, 1990; Dennett et al., 2002). Their significant participation to carbon fixation (through photosynthesis of their algal symbionts), and carbon export to the deep ocean (14,15) make them key players in oceanic waters. For instance, primary production of Collodaria can be more than four orders of magnitude greater than that in the same volume of the surrounding seawater (15). In addition to energy provided by their symbionts, *in situ* observations and culture experiments have also described collodarians as active predators feeding on a broad range of prey (e.g. copepods, ciliates, phytoplankton or bacteria) therefore contributing to oceanic food webs (16).

The ecological success of radiolarian photosymbioses in the ocean involving the microalga *Brandtodinium* must rely on the capacity of the partners to intertwine their metabolism and on the efficiency with which they exchange nutrients, such as carbon and the poorly available macronutrients (e.g. nitrogen) and trace metals (e.g. iron). However, the metabolism and physiology of the microalga *Brandtodinium* in symbiosis is not known, as well as the underpinning mechanisms of the host to accommodate this microalga. The study of this uncultured cell-to-cell symbiosis where partners cannot be separated calls for dedicated subcellular imaging methods that maintain the physical integrity of the host-symbiont association. Here, using multimodal subcellular imaging (3D electron microscopy, and mass spectrometry and X-ray fluorescence imaging), we investigated the cellular architecture and metal homeostasis of the microalga *Brandtodinium* in free-living and symbiotic stages, as well as the carbon and nitrogen flux and transfer between both partners.

## Results and Discussion

### Cellular architecture of the microalga *Brandtodinium* in free-living and symbiotic phases

The metabolism and bioenergetics features of a microalgal cell can be manifested by its cellular architecture organization, such as the arrangement and volume of energy-producing organelles and vacuoles (17). Using 2D and 3D electron microscopy (Transmission Electron Microscopy and Focused Ion Beam-Scanning Electron Microscopy - FIB-SEM, respectively), the cellular organization of the microalga *Brandtodinium* was investigated in the free-living (i.e. in culture) and symbiotic stage within the host Collodaria collected in surface waters of the Mediterranean Sea. Compared to free-living, the round-shaped microalgae have lost their thick theca in symbiosis (figure 1E-F). Of note, plastids, which were located around the cell periphery, occupied a 2-fold larger surface area of the cell in symbiosis (26 % of the cell surface ± 6; n= 20) compared to the free-living stage (12 ± 2 %; n = 41 cells) (figure 1G). In order to confirm and better understand this morphological change of the photosynthetic machinery, we conducted morphometric analyses using 3D electron microscopy (FIB-SEM) on four symbiotic *Brandtodinium* cells and three free-living cells maintained in culture. 3D reconstructions, which were based on >1000 aligned electron micrographs for each cell, showed that the volume of the microalgal cell was similar in both stages (333 ± 82 μm^3^, n = 3 and 305 ± 44 μm^3^, n = 4 in free-living and symbiosis, respectively). The plastid is a reticulated and thick network (figure 2 A,D). Morphometric analyses confirmed that the plastids were more voluminous in symbiosis (81 μm^3^± 12) compared to free-living (55 μm^3^ ± 10), and can occupy up to 31% of the cell volume (16.8 ± 1.5 % on average in free-living) (figure 2 G-H, Table S1). This expansion of the photosynthetic machinery was also found in the microalga *Phaeocystis* within acantharian hosts, suggesting that it may be a common morphological trait in planktonic photosymbiosis (17,18). The nutritional microhabitat provided by the host must be a favorable environment for expanding this plastid and optimize the capture of photons. Regarding the carbon fixation capability of *Brandtodinium*, 3D reconstructions unveiled various number of pyrenoids in individual cells (i.e. from one to four) in both life stages but in symbiosis, pyrenoid occupancy in the cell was slightly larger (2.4 ± 0.3%) than in free-living (1.7 ± 0.2%) (figure 2C, F). In addition, we observed a correlation between volumes of plastid and pyrenoid (R = 0.81, *p*= 0.028), suggesting an energetic coupling between the light-dependent and light-independent photosynthetic reactions (figure S1). In the multiply-stalked pyrenoids, we observed that thylakoid membranes (called tubules) penetrated like fingers the C-fixing organelle, which is covered by a thick starch sheath (figure 3A). These tubules are thought to be enriched in carbonic anhydrase in order to ensure high local concentration of CO_2_ for the Rubisco in the matrix (19,20).

**Figure 1:**
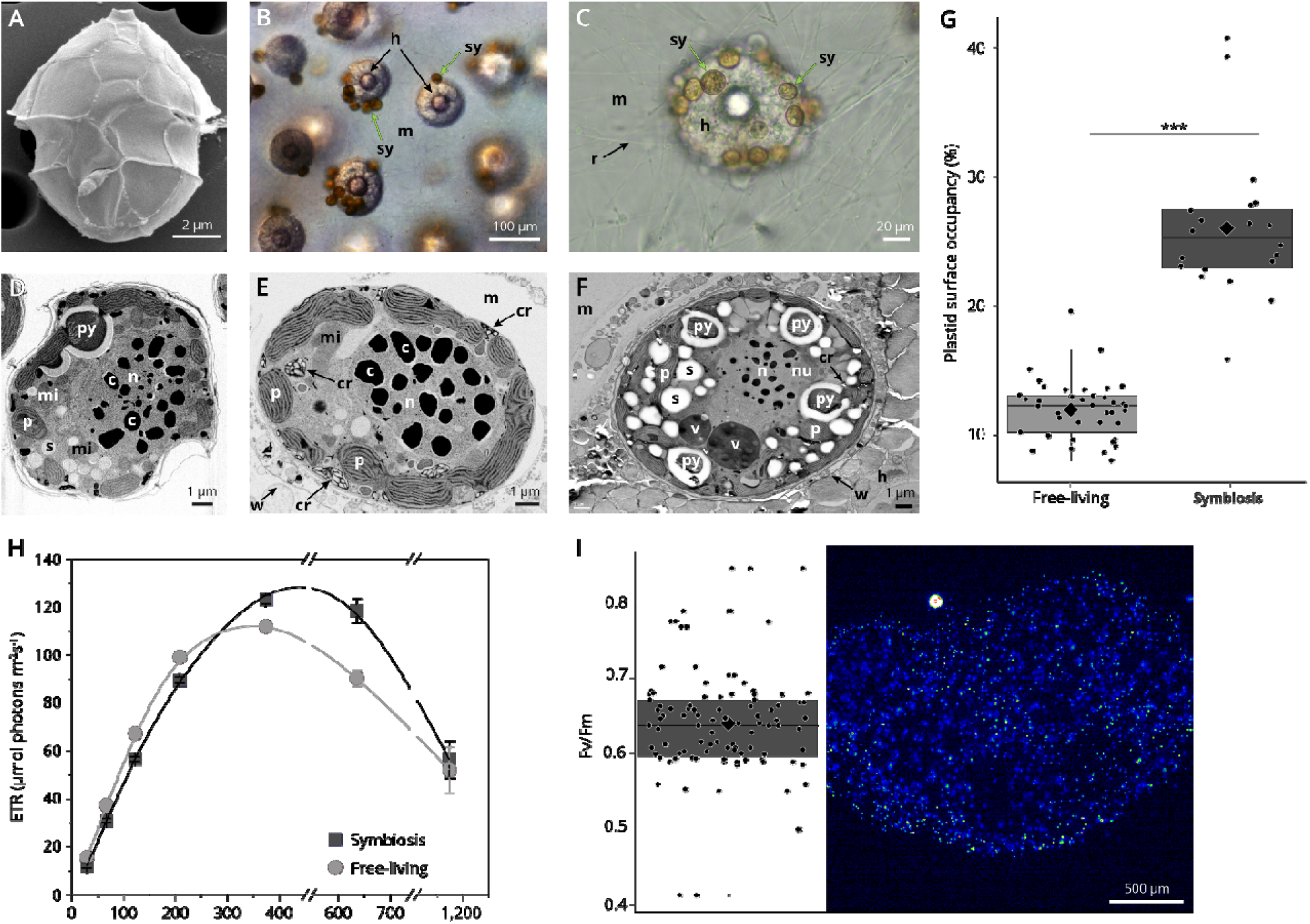
Ultrastructure and photophysiology of the microalga *Brandtodinium* in free-living and symbiotic stages within collodarians hosts. **A**: External morphology of *Brandtodinium* with its thecal plates revealed by Scanning Electron Microscopy. **B**: Colony of Collodaria observed with light microscopy, composed of several host cells (black arrow, h) and multiple symbiotic microalgae (Sy; green arrow) embedded in the gelatinous matrix (m). **C**: A zoom-in on a host cell surrounded by symbiotic microalgae and cytoplasmic strands (r; rhizopodia). **D**: Ultrastructure of free-living *Brandtodinium* revealed by Transmission Electron Microscopy (TEM). **E and F**: TEM micrographs showing the ultrastructure of symbiotic *Brandtodinium* within the Collodaria. **G**: Plastid surface area occupancy (% of the cell surface) in free-living and symbiotic stages of *Brandtodinium* calculated from TEM micrographs. Plastid surface area is statistically higher in symbiosis (t = 14.67, df = 24.75, p < 0.0001; N = 59; based on Cook’s distances and high leverage, two outlying data points were excluded from the analysis because of their high influence). **H**: Photosynthetic efficiency measured by the relative electron transfer rat (ETR) for free-living (light grey green circles; n= 4) and symbiotic (dark grey squares; n = 4) microalgae over a range of light intensities up to 1200 mmol photons m^−2^ s^−1^. **I**: F_v_/F_m_ values measured by a PAM microscope from each individual symbiotic microalgal cells within a colony. n: nucleus; c: condensed chromatin (chromosome), py: pyrenoid; p: plastid; s: starch grain; h: Host cell (Collodaria); m: matrix; mi: mitochondria; w: membranous envelope of the host; cr: nitrogen-rich crystals; r: rhizopodia.

**Figure 2:**
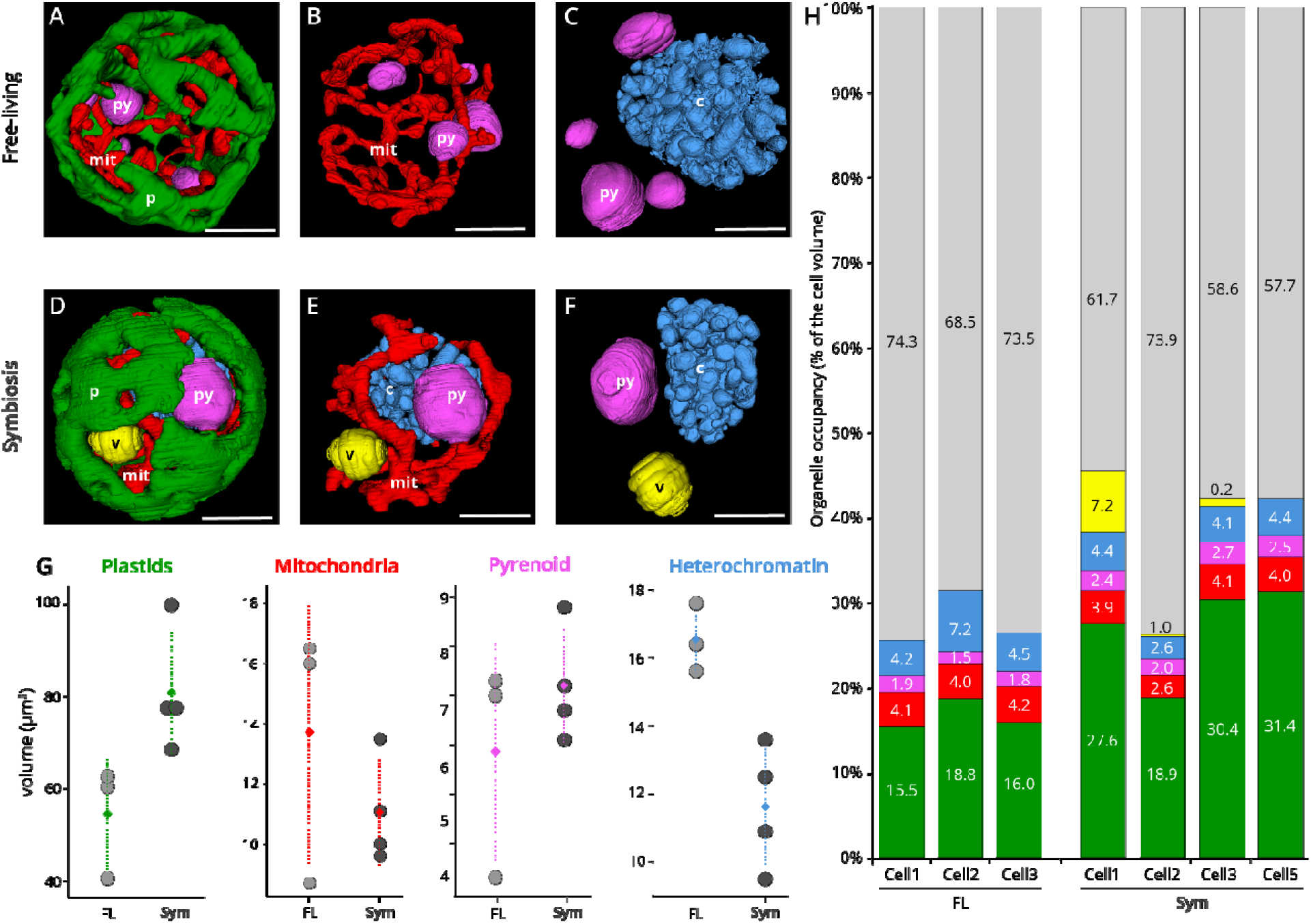
Cellular architecture in 3D and morphometrics of the microalga *Brandtodinium* in free-living and symbiotic stages revealed by FIB-SEM imaging. **A-C**: Topology of the plastid (green), mitochondria (red), pyrenoid (purple), and condensed chromatin (chromosomes, light blue) in a free-living microalgal cell. Scale bar: 3 μm. **D-F**: Topology of the plastid (green), mitochondria (red), pyrenoid (purple), vacuole (yellow) and condensed chromatin (chromosomes, light blue), in a symbiotic microalgal cell. Scale bar: 3 μm. **G**: Volume (μm^3^) of different organelles of the microalga *Brandtodinium* in free-living (n = 3) and symbiotic (n = 4) stages, such as plastids, mitochondria, pyrenoid and heterochromatin (chromosomes), calculated based on 3D reconstructions. **H**: Volume occupancy of organelles (% of the cell volume) in three free-living algal cells and four symbiotic algal cells.

**Figure 3:**
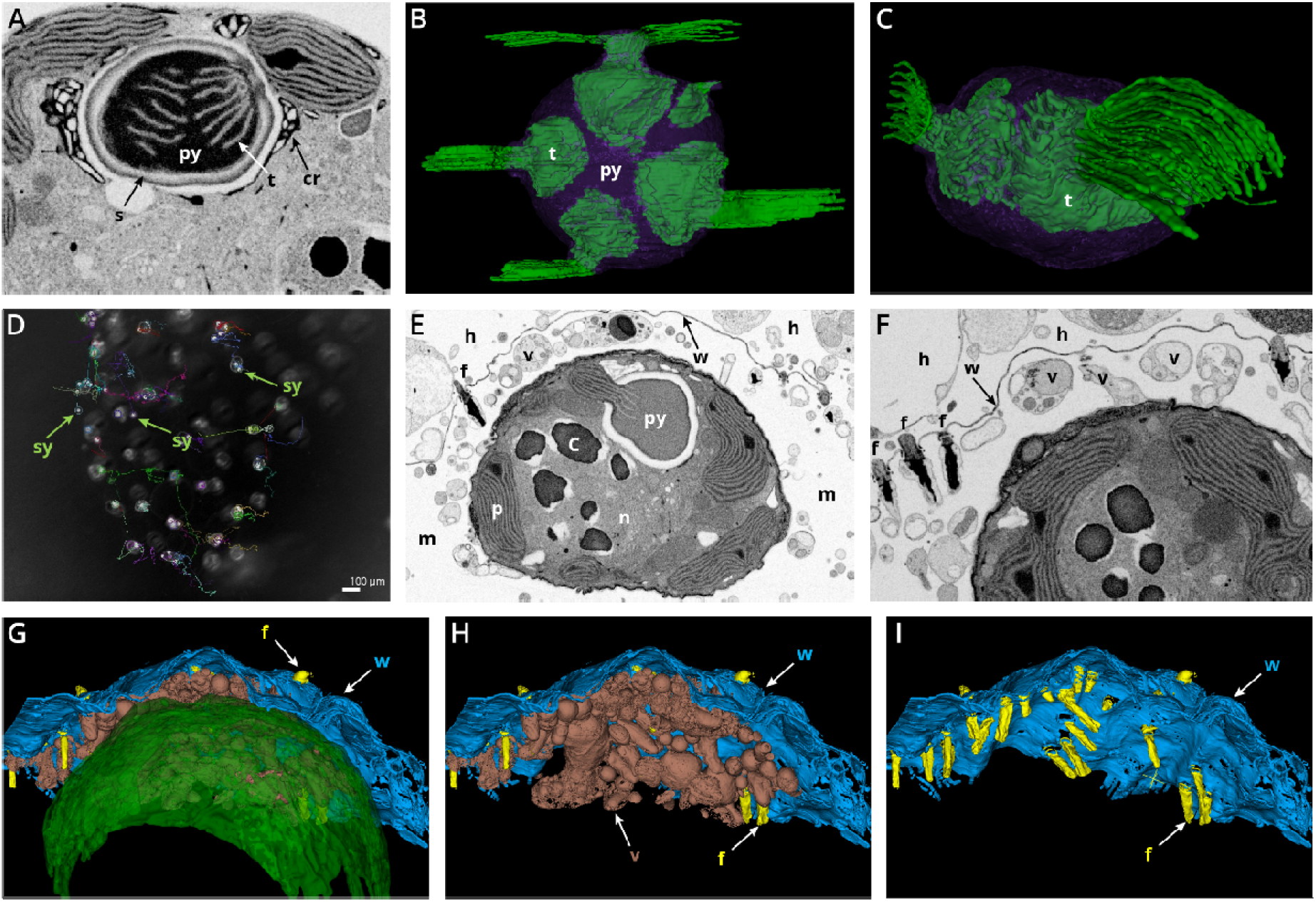
Pyrenoid architecture and integration of the symbiotic microalga in the host. **A-C**: Pyrenoid architecture of the microalga *Brandtodinium* visualized by TEM (A) and FIB-SEM reconstruction (B and C). Pyrenoid (purple) is surrounded by a starch sheath and penetrated by thylakoid membranes (t, in green) from different stalks. **D**: Snapshot image of a particle tracking video conducted on a light microscope (See video S1) where trajectories of symbiotic microalgae (Sy; indicated by a green arrow) during few hours are visible inside the colony (colored lines). Symbionts were not associated to a specific host cell but interacted with several host cells. **E-F**: EM micrographs showing the physical integration of a symbiotic microalga (*Brandtodinium*), which was surrounded by numerous vesicles (v) in the ectoplasm (matrix) and in close contact with the symbiont. **G-I**: 3D reconstruction after FIB-SEM of the host-symbiont integration with a focus on the numerous vesicles (v, brown structures) between the membranous envelope of the host (w, blue) and the symbiotic microalga (green). Piston-like structures (called fusules) were also observed at pore sites of the membranous envelope of the host (w). These organelles are known to inject host endoplasm into the matrix (=ectoplasm) as a vacuolated network (11). (n: nucleus; c: condensed chromatin (chromosome), py: pyrenoid; p: plastid; f: fusules; h: host cell (Collodaria); m: matrix; w: envelope of the host; cr: nitrogen-rich crystals; v: vacuoles in the matrix).

To test possible consequences of this plastid expansion on the photophysiology of *Brandtodinium*, we characterized photosynthetic parameters *in vivo,* based on chlorophyll fluorescence imaging. We found that the photosynthetic efficiency, measured through the electron transfer rate (ETR) parameter, which is related to carbon assimilation (21), was similar in free-living and symbiotic stages of the microalga in a light range from 20 to 400 μmol photons m^−2^ s^−1^, but significantly differed when light was further increased (being 1.3 times higher in symbiosis in the 400-1000 μmol photons m^−2^ s^−1^ range) (figure 1H). In addition, photoprotective response, evaluated though the Non-Photochemical Quenching parameter (NPQ) of photosystem II was lower in the symbiotic stage from 200 to 1200 μmol photons m^−2^ s^−1^ (being three times lower at 400 μmol photons m^−2^ s^−1^) (figure S2). Overall, these data suggest that the photosynthetic capacity is more efficient in symbiotic algae at high light (i.e. light intensity experienced in surface waters (22)), i.e. that absorbed photons are better used for carbon assimilation (ETR) and therefore less dissipated as heat. To test if photosynthesis is homogeneous among the symbiont population, photosynthetic capability was also measured on individual algal cells within a colony collected in surface waters (figure 1I). The average single-cell F_v_/F_m_ values was 0.64 ± 0.07 (n = 52 cells), consistent with earlier findings at the symbiont population level (0.58) (23). The variability observed among the symbiont population of the colony likely reflects different physiological states of symbiotic cells.

As for the respiratory machinery, the mitochondria formed a reticulated network located between the plastids and nucleus, in close proximity with plastids in some cellular regions (figure 2B, E). Total volume of mitochondria was not different between free-living (14 μm^3^ ± 4) and symbiotic cells (11 μm^3^ ± 2), occupying a constant 4% of the cell volume with similar topology in both stages (figure 2G, H). Based on this finding, it is tempting to propose that, although photosynthesis is modified, the respiratory activity of the microalgal cell is not substantially different in free-living and symbiosis. The nucleus of the microalga (also called the dinokaryon) was centrally located and contained typical condensed, rod-shaped chromosomes of different sizes and one or two nucleoli (24) (figure 2 C, F). The volume occupancy of the nucleolus in the cell was similar in both life stages (0.26 ± 0.4 % and 0.30 ± 0.07 % on average in free-living and symbiosis, respectively) (Table S1). But total volume of condensed chromatin tended to be slightly lower in symbiosis (12 μm^3^ ± 2; 3.9 ± 0.7% of occupancy) compared to that in free-living cells (17 μm^3^ ± 1; 5.3 ± 1.4% of occupancy) (figure 2G). The number of chromosomes including electron-dense chromatin structures in the nucleoplasm greatly varied from 44 to 88 in symbiotic cells and from 64 to 179 in free-living cells (figure S3). This reflects the dynamic level of chromatin compaction, which is known to vary across the life cycle of dinoflagellates (25). For instance, in G1 phase, there are numerous small structures protruding from the chromosomes toward the nucleoplasm (26), which could correspond to our observations in free-living cells (figure 2A). Although this needs to be confirmed with more cells, this observation may indicate that the symbiotic microalga is at a different life phase and may be related to the slower growth of symbiotic *Brandtodinium* as suggested by transcriptomics in a different host species and in corals (27,28). Yet, contrary to photosymbiosis in Acantharia, algal cells had the same volume as in free-living and were observed to divide within the host colony, so there is no complete inhibition of symbiont division (figure S4).

### Structural connectivity between the host and its symbiotic microalgae

The physical integration of symbiotic microalgae within the host was then investigated at the nanoscale to identify the possible routes of metabolic exchanges between cells. In Collodaria, microalgae were outside the host endoplasm, being embedded in the translucent gelatinous matrix of the colony (i.e. ectoplasm) (figure 3). Cytoplasmic strands (called rhizopodia) embraced microalgae and are known to control their distribution in the matrix (figure 1 (11)). Particle tracking from light microscopy time-lapse showed that microalgae can move in this matrix (up to hundreds of microns.min^−1^) and stop by different host cells for few minutes, likely for metabolic exchanges (figure 3D and Video S1). Of note, TEM and FIB-SEM reconstructions revealed a network of numerous vacuoles in the matrix located between the host cell and the symbiotic microalgae, sometimes physically associated to both cells (figure 3E-I). In 3D, some of these round-shaped vacuoles were connected each other and very likely ensure metabolic connection between the host and the microalgae. In addition, piston-like structures (called fusules) visible on the capsule membranous envelope of the host are known to inject host endoplasm into the matrix as a vacuolated network (11). Overall, Collodaria photosymbiosis is a dynamic system where host cells interact with several microalgae within the “greenhouse-like” colony through time via vesicles, raising questions about the nutrient flow between cells and more generally the metabolic connection.

### Subcellular visualization of the incorporation and allocation of ^13^C and ^15^N

To better understand the metabolic crosstalk between the host and the microalgae and more specifically the exchanges and allocation of carbon and nitrogen, we incubated collodarians in ^13^C- bicarbonate and ^15^N-ammonium for three-hour light period in filtered natural seawater, cryo-fixed at high pressure and prepared them for nanoSIMS (Nanoscale Secondary Ion Mass Spectrometry) (See methods). Five different microalgae from two distinct collodarian colonies were analyzed. NanoSIMS analyses revealed increase in ^13^C fraction (% of ^13^C out of total C) mainly in multiple starch grains distributed in the microalgal cytoplasm (21 ± 2 at%, n = 38) and in the extra-plastidial starch sheaths surrounding the pyrenoids (19 ± 1 at%, n = 12) (figure 4, Table S2). With the detected increase in heavy isotope fractions, the relative carbon assimilation Ka of 25 ± 4 at% in starch grains and 23 ± 2 at% in starch sheaths were calculated (see methods for more details and (29)). Lower relative carbon assimilation (Ka) was revealed from starch of the second collodarian colony (10 ± 3 at% and 9 ± 2 at% in cytoplasmic grains and sheaths, respectively). Starch grains were observed both in free-living and symbiotic stages by TEM observations (figure 1), and can represent a volume of ca. 5% of the cell as calculated from FIB-SEM reconstructions (figure 4, Table S1). They were scattered in the cytoplasm and adjacent to plastids and pyrenoids. ^13^C-labelled starch granules were also observed in another planktonic photosymbiosis in foraminifera involving the dinoflagellate *Pelagodinium beii* (30). The observed assimilation activity pattern in photosymbiosis confirms that the newly-fixed carbon is rapidly stored by the microalgae rather than incorporated into the algal biomass.

**Figure 4:**
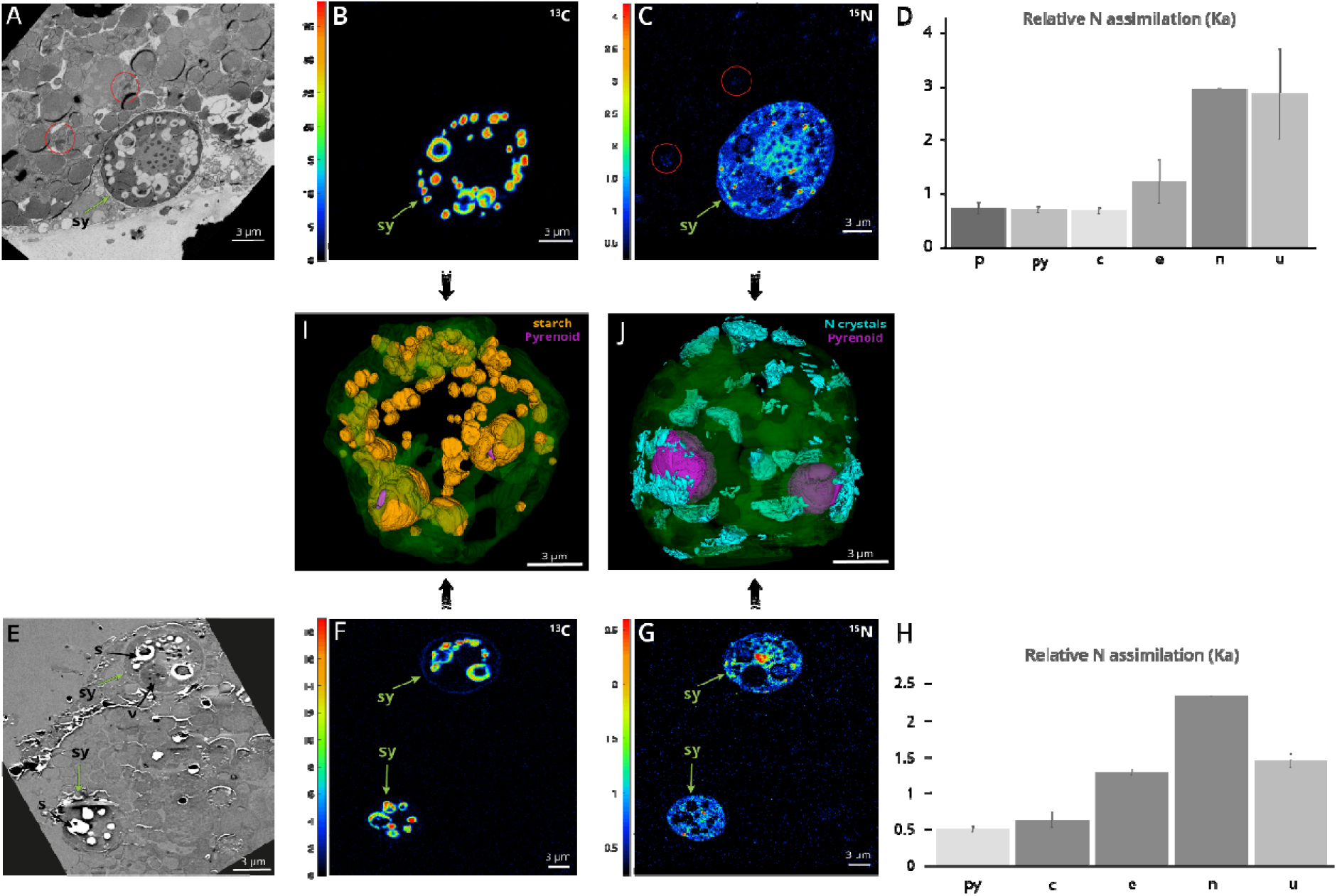
Carbon and nitrogen flux in the symbiotic cells as revealed by correlated TEM-nanoSIMS and ^13^C-bicarbonate and ^15^N-ammonium labeling. **A and E**: TEM micrographs of the cellular areas analyzed by nanoSIMS, showing three symbiotic microalgae (Sy; indicated by a green arrow) and their host cell from two different colonies. Host Golgi are surrounded by a red circle. **B and F**: NanoSIMS mapping showing relative carbon assimilation (Ka) following ^13^C-bicarbonate 3h-incubation in starch granules and starch surrounding the pyrenoids. **C and G**: NanoSIMS mapping showing the relative nitrogen assimilation (Ka) following ^15^N-ammonium 3h-incubation in symbiotic microalgae, more particularly in purine crystals (red hotspots) and in the nucleolus. The latter compartment corresponds to the red hotspot in the middle of the upper cell in panel G. Nitrogen assimilation was also detected in the Golgi apparatus of the host cell, highlighted by the red circles in C (see also figure S5). **D and H**: Averaged relative N assimilation (Ka) measured by nanoSIMS in different organelles of the symbiotic microalgae from two different colonies (A8 and A7) (n= 3 cells in D; n = 2 cells in H). **I**: 3D reconstruction after FIB-SEM imaging of the starch (orange) that assimilated carbon. In the algal cytoplasm, starch was contained in grains and also surrounding the pyrenoids (purple) and closely associated to the plastid (green, transparent). **J:** 3D reconstruction after FIB-SEM imaging of N-rich purine crystals (light blue) that were localized at the periphery of the cell in close association with the pyrenoids (purple). py: pyrenoid; c: condensed chromatin; e: euchromatin+nucleoplasm; n: nucleolus; u: purine crystals.

In the host cell, after three hours of incubation, low ^13^C assimilated fraction was detected and only in the secretory vesicles of multiple Golgi apparatus (1.33 ± 0.05 at%, corresponding to relative carbon assimilation (Ka) of 1.64 ± 0.12 at% (figure 4, figure S5). This organelle is known to serve as a carbohydrate factory to process oligosaccharide side chains on glycoproteins and glycolipids (Mellman & Simons 1992). Once translocated from the symbiont to the host, we therefore presume that ^13^C is incorporated into host biomass such as proteins and lipids through the Golgi apparatus (31). Overall, carbon assimilation pattern into the host seems to be different in Collodaria-*Brandtodinium* symbiosis compared to other dinoflagellate photosymbioses in corals (Symbiodiniaceae) and foraminifera (*Pelagodinium)* (30,32). In these symbioses, ^13^C-enrichment were first observed in the host lipid droplets after 2 hours, and later elsewhere in the host cell. In this study, no putative lipid droplets were observed in collodarian hosts in different sections observed with electron microscopy and may explain the ^13^C assimilation differences compared to other photosymbioses. We may also consider that most of the carbon fixed and stored by the symbionts during the day could be transferred overnight to its host, so explaining low ^13^C fraction in the host in this study. One analytical aspect to take into account is that NanoSIMS provides a 2D information so depending on the cellular sections, some important organelles can be missed in the imaging areas, such as lipid droplets or vacuoles. In the matrix of the colony, the vacuoles that we hypothesize to be a route for metabolic exchanges were not showing carbon assimilation activity. This may be explained by the fact that soluble photosynthates (sugars), which are transferred to hosts in photosymbioses (28), are lost during sample preparation, so could not be visualized with nanoSIMS (33).

The subcellular distribution of nitrogen (^12^C^14^N^−^/^12^C_2_^−^, figure S6), which is limiting in the ocean and mainly contained in proteins and amino acids (34,35), showed that *Brandtodinium* symbionts contained a significant amount of nitrogen (N) compared to the host (Table S3). In free-living and symbiotic *Brandtodinium* cells, N was mainly contained in similar amount in plastids, pyrenoids and nucleus. The N-rich photosynthetic machinery is due to the high concentration of photosystems and pigments in the plastid and the CO_2_-fixing Rubisco in the pyrenoid. In the nucleus, the chromosomes and the nucleolus were particularly rich in N, likely representing ribosomes and proteins associated to the chromatin (figure S6). Overall, subcellular mapping of nitrogen pinpointed metabolic needs of the symbiotic microalgae and showed that the host must provide a significant amount of N to support the functioning of the nucleus and photosynthetic machinery. Collodaria are known to be active predators (16), therefore representing a source of nitrogen for their symbionts. In addition, like other hosts, we presume that collodarians can uptake inorganic nitrogen such as ammonium and transfer it to its symbionts, but this remains unknown.

In order to reveal the first steps of N flux between the two partners, symbiotic associations were incubated with ^15^N-labelled ammonium for 3 hours. High ^15^N assimilation fraction was measured in small hotspots in the algal cell from both colonies: up to 3.1 ± 0.8 at% which correspond to 2.9 ± 0.9 at% of relative N assimilation (Ka). These ^15^N hotspots correspond to crystalline inclusions as seen by electron microscopy (figures 1E, 3A, 4J). FIB-SEM reconstruction showed that, in the symbiotic stage, N-rich crystals were located at the cell periphery and can represent ca. 2% of the algal cell volume (figure 4). Based on their morphologies in EM, these crystals likely correspond to the purines guanine or uric acid, which are known to be nitrogenous reserves in microalgae including the symbiotic *Symbiodiniaceae* in corals (36–38). Note that we did not observe these crystals in the free-living microalgae, which may be explained by the fact that they grow in a N-replete culture medium so do not need to store N. These crystals showed close physical contacts with the plastids and pyrenoids (figure 3A), indicating that these organelles may either produce or benefit from this N reserve. The presence of purine crystals would ensure N source in N-depleted waters or when prey feeding by the host is restricted. Future studies are required to better understand the dynamics of this N reserve in symbiotic microalgae over 24 hours and in different trophic conditions (e.g. hosts in starved or N-replete conditions).

^15^N assimilated fraction was also important in the nucleolus of the algal nucleus (2.9 at% ± 0.3, or 2.7 at% ± 0.3 of relative assimilation, Ka). This suggests that N is preferentially utilized and allocated for the synthesis of ribosomes inside the N-rich nuclear apparatus. Euchromatin was also ^15^N-enriched (1.6 at% Ka) and then similar Ka values were found in the condensed chromatin, plastid and pyrenoid (ca 0.6 at% Ka) (figure 4). In addition to N, subcellular mapping of phosphorous (^31^P/^12^C_2_) showed that the nucleus of *Brandtodinium* is the organelle that contains the more P in the algal cell, more specifically in the condensed chromatin with up to ca. 10 times more than in other organelles (figure S6, Table S3).

In the host cell, low ^15^N fraction was found (0.76 ± 0.05 at%; ka: 0.40 ± 0.05 at%) in the Golgi apparatus, including cisternae and secretory vesicles (figure 4 and figure S5). Whether this ^15^N originates from a direct ammonium assimilation by the host cell or from N-exchanges with the symbiotic dinoflagellates cannot be resolved in this study and would require shorter incubations (e.g. less than 1 h). Overall, ^15^N-assimilation in the collodarian host cell seems to be slower compared to other protistan photosymbioses. For example, in foraminiferal symbiosis, ^15^N-enrichment in the host cell was detected in specific structures already after one hour of incubation with ^15^N-ammonium and rapidly spread to the entire host cell (39).

### Subcellular distribution of sulfur

Sulfur plays a pivotal role as protein constituent (e.g. metalloproteins) and as antioxidant protection with the production of several compounds, such as dimethylsulfoniopropionate (DMSP), which contribute to the global sulfur cycle of the ocean (40,41). Quantification using Synchrotron X-Ray Fluorescence (S-XRF) imaging revealed that symbiotic microalgae contained around two times more sulfur than their host Collodaria (figure 5, Table S4). Similar subcellular distribution and concentrations were found between symbiotic (2400 ppm ± 600; n = 13) and free-living (2100 ppm ± 400; n = 33) *Brandtodinium* cells. NanoSIMS showed that sulfur was mostly located in the condensed chromatin, and then about two times less in pyrenoids and plastids in descending order (figure 5, Table S4). This is in contrast to what is known in other microalgae where most sulfur is contained in the photosynthetic machinery (8,40). In addition, in the symbiotic microalgae, nanoSIMS revealed a large vacuole of 2.7 μm in size, which contained much more sulfur than chromatin (figures 1, 2 and 5). FIB-SEM reconstruction showed that this vacuole can represent up to 7% of the algal cell volume (figure 2). This vacuole did not assimilate ^13^C and ^15^N and contained no phosphorous (figure 6 and S6). It is known that *Brandtodinium* can produce 100-fold higher DMSP compared to the free-living form (Gutierrez-Rodriguez et al. 2017). Therefore, this S-rich vacuole could contain DMSP and DMS as also observed in Symbiodiniaceae in culture (43) or other S-rich molecules involved in the algal metabolism. Of note, in the matrix of Collodaria, sulfur was also highly concentrated (5400 ppm ± 800) in the numerous extracellular vacuoles reconstructed in 3D above (figures 3 and 5, Table S4). These vacuoles contained four and two times more sulfur than in the host endoplasm and symbiont, respectively. This would indicate that sulfur-containing molecules and/or proteins could be exchanged between both partners, thus stressing the role of the sulfur metabolism in photosymbiosis. Future analyses using complementary mass spectrometry imaging (e.g. ToF-SIMS) should be conducted to identify the S compounds in symbiotic cells at relatively high resolution (< 1μm) (43,44).

**Figure 5:**
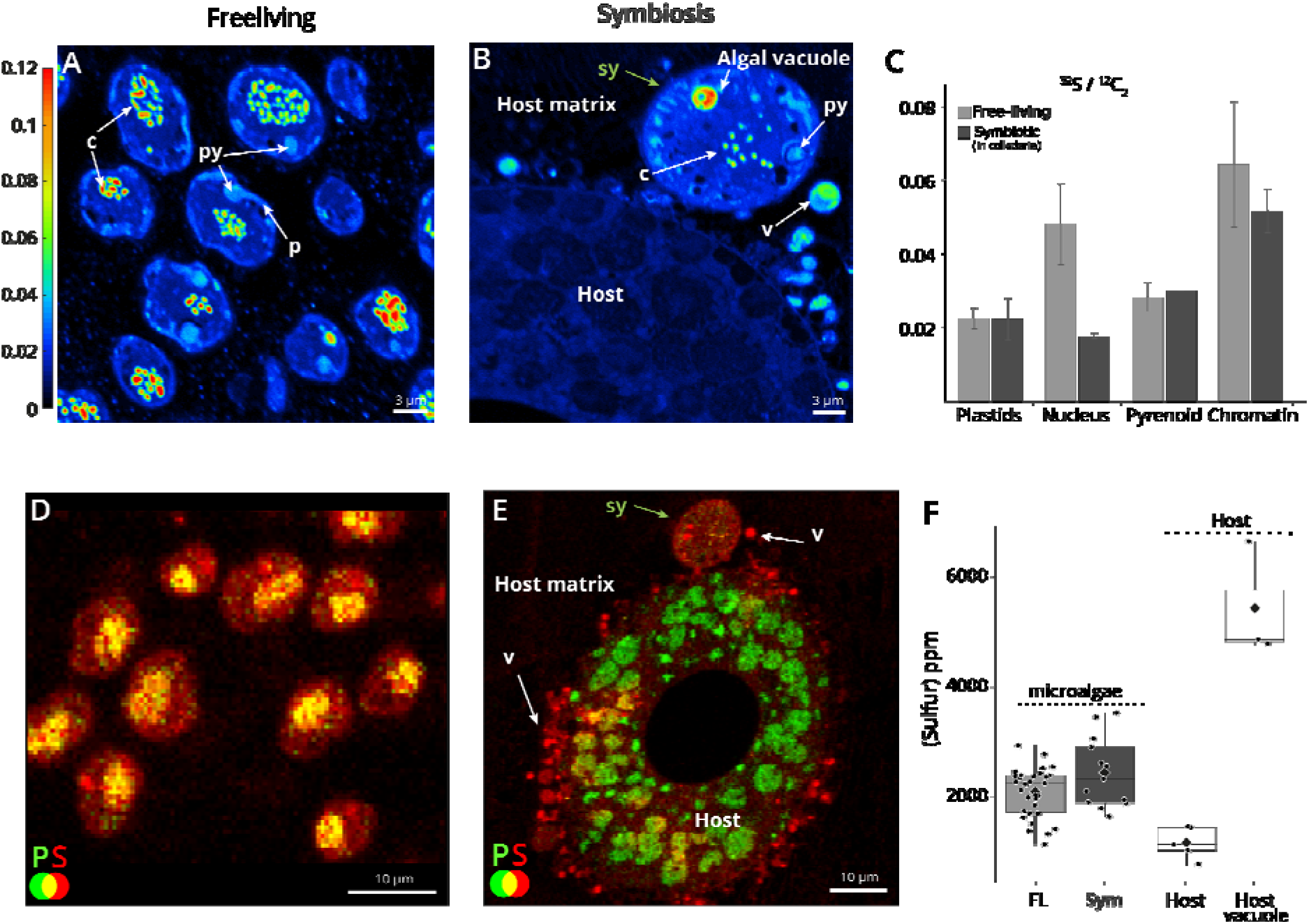
Subcellular distribution of sulfur in the collodaria photosymbiosis unveiled by nanoSIMS and Synchrotron X-Ray Fluorescence (S-XRF). **A and B**: Sulfur (^32^S/^12^C_2_) mapping in free-living (A) and symbiotic (B) microalgae unveiled by nanoSIMS. Sulfur was mainly localized in chromatin of microalgae, but also in a large vacuole in the symbiotic cell (green arrow in B). Sulfur was also concentrated in vacuoles of the matrix between the host cell and the symbiotic microalga. Scale size: 3 μm. **C**: Normalized sulfur content (^32^S/^12^C_2_) measured by nanoSIMS in different organelles (plastids, nucleus, pyrenoid and chromatin) of free-living (light grey) and symbiotic (dark grey) microalgae. (Table S3) **D and E**: Sulfur (red) and phosphorous (green) distribution unveiled by S-XRF imaging showing high concentration of sulfur in microalgae (plastids, chromatin) and the vacuoles of the matrix. The co-localization of S and P is indicated by the yellow color (nucleus of the microalga in D). **F**: Sulfur concentration (ppm) measured by S-XRF in the free-living (n = 33) and symbiotic *Brandtodinium* (n =13) cells (dark and light grey, respectively), as well as in the host cell (n = 5) and the vacuoles of the matrix. (n: nucleus; c: condensed chromatin; py: pyrenoid; p: plastid; h: host cell (Collodaria); m: matrix; v: vacuoles in the matrix).

**Figure 6:**
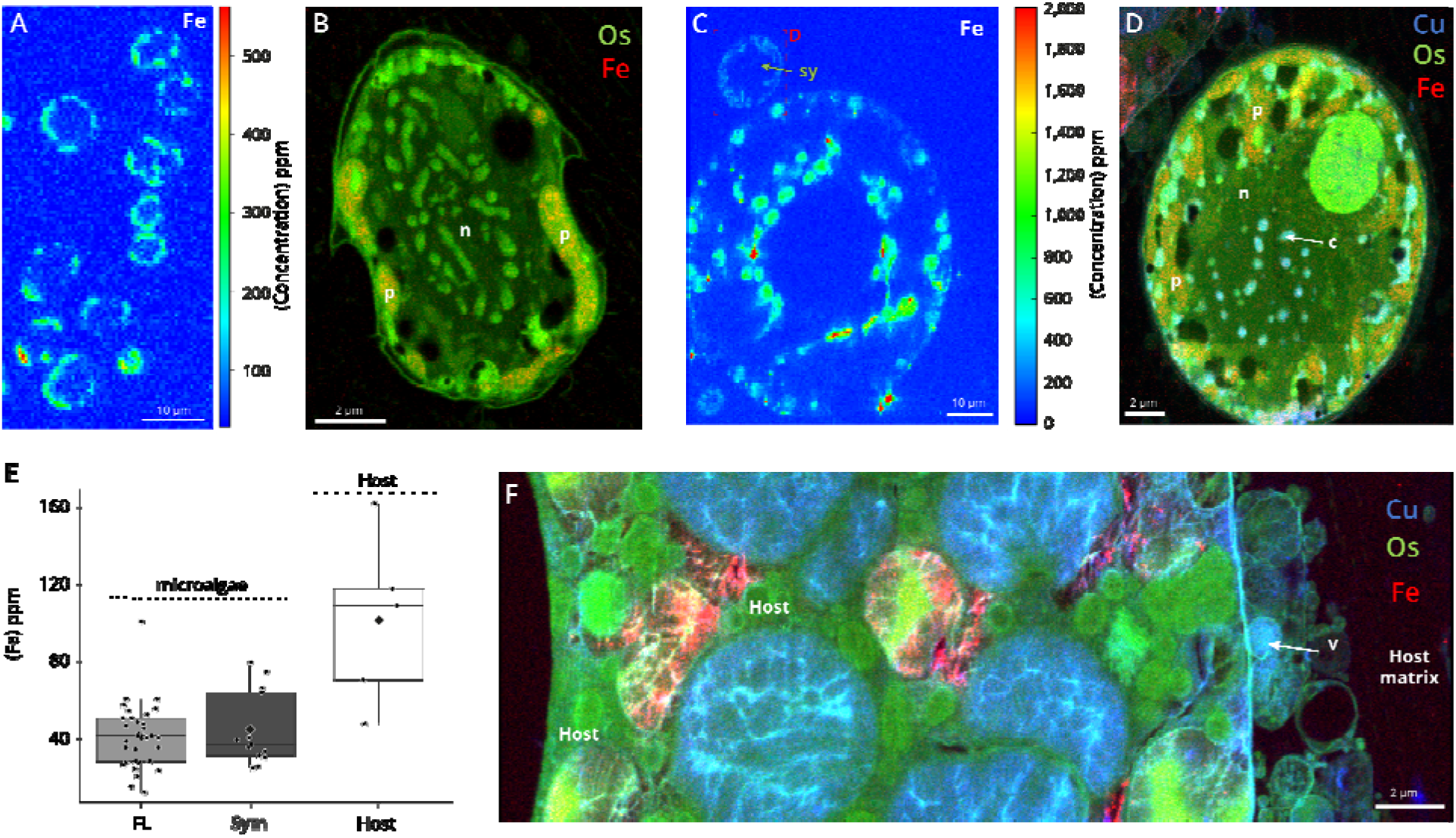
Concentration and subcellular distribution of the trace metals iron (Fe) and copper (Cu) in microalgal and host cells unveiled by Synchrotron X-ray Fluorescence. **A, C**: Subcellular quantitative mapping of iron (Fe) showing its concentration (ppm) in free-living microalgae (A), and in a host and its symbiotic microalga (C) (500 nm xy resolution and at 7.3 keV). Scale bar: 10 μm. **B, D and F**: High-resolution mapping (50 nm xy resolution and at 17.5 keV) of osmium (Os, green, providing ultrastructure information), iron (Fe, red) and copper (Cu, blue) in free-living (B) and symbiotic (D) microalgae, and in the host cell (F) extracted from XRF hyperspectral images. Scale bars: 2 μm. Fe was mainly concentrated in plastids and host vacuoles. Note that Cu was only detected in symbiotic microalgae in the condensed chromatin (c) and cytoplasmic vacuoles, and was below the detection limit in free-living cells (See also figure S8). **E**: Intracellular iron concentration (ppm) measured by S-XRF in the free-living (FL; n = 33) and symbiotic *Brandtodinium* (Symb, n =13) cells (dark and light grey, respectively), a well as in the host cell (n = 5). (n: nucleus; c: condensed chromatin; py: pyrenoid; p: plastid; h: host cell (Collodaria); m: matrix; v: vacuoles in the matrix).

### Subcellular mapping of metals

Trace metals are essential for diverse biochemical functions, such as photosynthesis, antioxidant and photo-protection (45,46). The ecological success of photosymbiosis in oceanic waters must therefore rely on efficient mechanisms to uptake and regulate the exchange of these poorly available elements. More particularly, iron (Fe) has been shown to be pivotal for the physiology and health of coral photosymbioses (47). Yet, trace metal content in planktonic photosymbioses remain largely unexplored, and their visualization and quantification in cells can provide information not only on the functioning of photosymbiosis, but also of the impact in trace element cycling, especially because collodarians are important agents in vertical flux (14).

In collodarian colonies (including the symbiotic microalgae), metallome analyses showed than Mn, Fe, Zn and Ni were the main trace metals (intracellular metal quotas normalized against phosphorus as the biomass indicator: ranging from 10 to 30 mmol/mol P), while Co and Cu were measured in relatively lower concentration (from 0.1 to 20 mmol/mol P) (figure S7; Table S5). We can note that Fe quota is about three times more than that of zooplankton and phytoplankton communities (4.6, 5.2, 4.9, 4.7 mmol Fe/mol P) sampled in different environmental settings (45). In other words, one collodarian colony of several mm long can contain 100-300 ng of Fe.

Two S-XRF beam lines were used to visualize and quantify Fe in cells at 500 nm and 50 nm spatial resolution of with a beam energy of 7.3 and 17.5 keV, respectively. In the host, Fe concentration tended to be two-fold higher (100 ± 40 ppm; n = 5) than Fe in their symbiotic microalgae (45 ± 18 ppm; n = 13) (figure 6). More specifically, the host cytoplasm exhibited round structures with high concentration of Fe (ca 400 ppm). In microalgae, Fe was mainly concentrated in plastids and its concentration tended to be similar between free-living (42 ± 16 ppm; n = 33) and symbiotic cells (figure 6B). Yet, the comparison is difficult between free-living and symbiotic microalgae, which grow in Fe-replete culture medium (5.85 μM) and in the host cell, respectively. Nevertheless, the fact that microalgae seem to have similar Fe concentration may indicate that the host could provide a Fe-replete microhabitat for its symbionts. It is known that the dinoflagellates Symbiodiniaceae increase their intracellular Fe concentration as the external Fe concentration increases (47,48).

Using high-resolution and high-energy S-XRF beam line, copper (Cu) was also detected and its distribution mapped in cells. Of note, Cu was only detected in the symbiotic stage of the microalga but not in free-living (below the detection level) (figure 6 and figure S8). More specifically, Cu was highly concentrated (6900 ppm) in the condensed DNA chromatin and vacuoles physically associated to plastids. In the host, Cu was present in large round structures and in some small vacuoles of the matrix (1900 ppm) (figure 6), which, like sulfur, may reflect the exchange of this metal between the host and the symbionts. The increase of Cu concentration in symbiotic microalgae represents a symbiotic fingerprint and stresses the influence of the host on their metal homeostasis. It is known that Cu is needed in the electron transport chain of the plastid (plastocyanin) and mitochondria (cytochrome c oxidase) and is a key cofactor in many proteins (49), but this study cannot resolve the function of this metal in the symbiosis. Nevertheless, Cu is known to be toxic at very low concentration (46), induce ROS production and DNA damage and is implicated in various neurological disorders, implying that its exchange between symbiotic partners should be highly regulated. Overall, the subcellular mapping of trace metals calls for future studies on the role of these metals in the symbiotic interaction. Because collodaria are significant contributors to the vertical flux in the ocean (13), these results also stress the role of photosymbiosis in the biogeochemical cycles of trace metals.

## Conclusions

The aim of this study was to improve our understanding on the functioning and metabolism of the uncultivated yet ubiquitous oceanic photosymbiosis between Collodaria and its symbiotic microalgae *Brandtodinium* using multimodal subcellular imaging. We showed that the volume occupancy of the plastid and C-fixing pyrenoid of the microalga increased in symbiosis and was accompanied by a higher photosynthetic performance at high light. By contrast, the topology and volume of the mitochondria was similar, suggesting similar respiration rate in free-living and symbiosis. The enlargement of the photosynthetic apparatus seems to be a morphological trait in planktonic photosymbiosis as it was also found in other radiolarians (Acantharia) living with the microalga *Phaeocystis* (8,9). Yet, the expansion of the photosynthetic machinery observed here in *Brandtodinum* is less substantial than in *Phaeocystis*, which can increase by 100-fold the volume of total plastids (9). This may be explained by the level of integration and growth of the microalgal cell within its host. In Collodaria, symbiotic microalgae are outside the host endoplasm in the gelatinous matrix, can divide (possibly at lower rate than free-living cells in culture) and they are not associated to a single host cell but can freely move within the colony interacting with several host cells over time. By contrast, the level of integration and host control is higher in Acantharia since the microalga *Phaeocystis* is constrained within the endoplasm of its host and its cell division is blocked but plastids continue to divide producing a highly voluminous algal cell. This indicates that radiolarian hosts have different strategies to control and accommodate their symbiotic microalgae.

Based on 3D electron microscopy and chemical imaging, we suggest that the metabolic connection in Collodaria between partners occurs through numerous extracellular vacuoles in the matrix, which contained nutrients, such as nitrogen, sulfur and copper. These nutrients are therefore important for the functioning of the symbiosis and calls for future analyses to identify the underlying metabolites. We demonstrated that carbon and nitrogen was stored by the symbiotic microalgae during the day and transferred in low amount in the Golgi apparatus of the host. This transfer seems to be less efficient than other planktonic and benthic photosymbioses (37,39), presumably due to this loose interaction between the symbionts and the host cells. Dynamic nutrient exchange at different time points during day and night periods should be carried out in future studies to fully understand the metabolic connection and flux of nutrients in this ecologically-successful photosymbiosis. Overall, the visualization of the morphology and metabolism of this symbiosis at the nanoscale further stresses the ecological importance of Collodaria in the biogeochemical cycles of carbon, sulfur and metals in the ocean.

## Author Contributions

J.D. conceived and designed research, and drafted the manuscript; J.D., G.V., C.L., H.S., S.R., G.F. participated in data analysis, and helped draft and critically revised the manuscript. N.M., B.G., F.C., S.M., R.T., N.S., Y.S. contributed to data acquisition, culture collection, sample preparation and helped draft the manuscript. All authors gave final approval for publication and agree to be held accountable for the work performed therein.

## Acknowledgements

This research was supported by the Dept. of Isotope Biogeochemistry, Centre for Chemical Microscopy (ProVIS), Helmholtz Centre for Environmental Research (UFZ). We thank Estelle Bigeard and Fabien Lombard for sample collection and the institutes the Laboratoire d’Océanographie de Villefranche-sur-Mer (LOV) and the marine service crew. This research is also supported by EMBRC-France, whose French state funds are managed by the ANR within the Investments of the Future program under reference ANR-10-INBS-02. This work used the platforms of the Grenoble Instruct centre (ISBG; UMS 3518 CNRS-CEA-UJF-EMBL) with support from FRISBI (ANR-10-INSB-05-02) and GRAL (ANR-10-LABX-49-01) within the Grenoble Partnership for Structural Biology (PSB). The electron microscope facility is supported by the Rhône-Alpes Region, the Fondation Recherche Medicale (FRM), the fonds FEDER, the Centre National de la Recherche Scientifique (CNRS), the CEA, the University of Grenoble, EMBL, and the GIS-Infrastrutures en Biologie Sante et Agronomie (IBISA). The FIB-SEM work was done in collaboration with the electron microscopy core facility at EMBL Heidelberg. The authors acknowledge the support and the use of resources of Instruct, a Landmark ESFRI project. The authors acknowledge the ESRF for providing beamtime for this project (LS-2625 and LS-2857 on ID16B-NA, in-house beamtime on ID21), and Marine Cotte and Hiram Castillo-Michel for their help in the experiments. We thank Louise Soegaard Jensen for access and help with SEM imaging, and Abby Ren, Stephane Escrig and Anders Meibom for NanoSIMS access and assistance. Authors are grateful to Dr. Lubos Polerecky (Utrecht University) for the continuous development of the LANS software, technical assistance and prompt implementation of software features.

## Material and Methods

### Sampling and isolation of Collodaria and microalgae

Colonies of collodarians (radiolarians) were gently collected at the subsurface in the Mediterranean Sea (Villefranche-sur-Mer, France) using a plankton net of 150 μm in mesh size. Live collodarians were then isolated through the binocular, maintained in filtered seawater and controlled light and temperature conditions, and cryofixed with high pressure freezing to preserve the native ultrastructure and chemistry of cells (50–52). In parallel, the microalgae (the dinoflagellate *Brandtodinium nutricula* RCC 3387 (10)) grown in culture medium K2 (http://roscoff-culture-collection.org/culture-media) at 20°C were also cryofixed in the exponential phase in the same conditions. Time lapse video using an inverted microscope was conducted on live colonies upon collection during several hours with image capture every 20 seconds. Trajectory of symbiotic microalgae over time was reconstructed with the plugin Particle Tracker for Fiji (53).

### Photophysiology

Photosynthetic activity was imaged with a fluorescence imaging setup described in (8). The photosynthetic electron transfer rate, ETR, was calculated as the product of the light intensity times the photochemical yield in the light: PFD × (F_m_’−F)/F_m_’), where F and F_m_’ are the steady-state and maximum fluorescence intensities in light-acclimated cells, respectively, and PFD (photosynthetic flux density) is the light irradiance in μmol quanta *m^−2^ s^−1^. The light intensity was increased stepwise from 29 to 1200 μmol quanta × m^−2^ s^−1^. The photoprotective responses were evaluated by measuring the non-photochemical quenching of fluorescence (NPQ,(54)) using the fluorescence setup described above. The NPQ was calculated as 1-(F_m_’/F_m_). The F_v_/F_m_ parameter (maximum potential quantum efficiency of Photosystem II) was assessed as F_m_-F_o_/F_m_ where F_m_ and F_0_ are the maximum and minimum fluorescence intensities. This parameter was measured in single cells using a homemade fluorescence imaging setup. The system uses a high sensitivity camera (Orca Flash 4.0 LT, Hamamatsu, Japan) equipped with a near infrared long pass filter (RG 695 Schott, Germany), mounted on an optical microscope (CKX 53 Olympus, Japan). Fluorescence was excited with pulses (duration 260 μs) provided by a blue LED (λ= 470 nm ± 12 nm) to induce the F_o_ level. Green LEDs (λ = 520 nm ± 20 nm) were used to generate short saturating pulses (intensity 3000 μmol photons m^−2^ s^−1^, duration 250ms) to induced F_m_, which was also measured with the blue pulses, given 10 μs after the saturating light was switched off. The blue and green LEDS were mounted on an array located on the upper part of the microscope, i.e. opposite to the objectives used for detection. We used a 20X (NA= 0.45) objective to scan the slits in the transmission mode.

### Sample preparation for electron microscopy and chemical imaging

Rapid freezing methods are universally accepted as superior to chemical fixation in preserving cell ultrastructure and chemistry (52). Symbiotic collodarians (host and algal symbionts) and free-living microalgae were therefore cryo-fixed as in (8), using high-pressure freezing (HPM100, Leica) where cells were subjected to a pressure of 210 MPa at −196°C for 30 ms, followed by freeze-substitution (EM ASF2, Leica). For the freeze substitution (FS), a mixture 2% (w/v) osmium tetroxide and 0.5% (w/v) uranyl acetate in dried acetone was used. The FS machine was programmed as follows: 60-80 h at −90°C, heating rate of 2°C h^−^1 to −60°C (15h), 10-12h at −60°C C, heating rate of 2°C h^−^1 to −30°C (15h), 10-12 h at −30°C. Samples were then washed in acetone four times for 20 minutes at −30°C and embedded in anhydrous araldite. Without accelerator, a graded resin/acetone (v/v) series was used (30, 50 and 70% resin) with each step lasting 2h at increased temperature: 30% resin/acetone bath from −30°C to −10C°, 50% resin/acetone bath from −10°C to 10°C, 70% resin/acetone bath from 10°C to 20°C. Samples were then placed in 100% resin without accelerator for 8-10 h and in 100% resin with accelerator (BDMA) for 8 h at room temperature. Prior to ultra-sectioning, symbiotic cells were observed in the resin block to define the trimming region and the z-position of cells in the block. Trimming around the targeted cells was performed with razor blades and the EM Trimming Leica machine. Thin sections (200-400 nm thick) were then obtained using an ultramicrotome (Leica EM) with ultra-diamond knife (Diatom) and placed on 10-mm arsenic-doped silicon wafers for NanoSIMS, and on Si_3_N_4_ membrane windows for synchrotron X-rays fluorescence. Adjacent sections of 60-80 nm thick were also obtained for TEM analysis.

### FIB-SEM acquisition and analysis

For FIB-SEM, upon high pressure freezing, the cocktail of the freeze substitution contained 2% (w/v) osmium tetroxide and 0.5% (w/v) uranyl acetate in dried acetone as in (9,17). The cells were washed four times in anhydrous acetone for 15 min each at −30 °C and gradually embedded in anhydrous araldite resin. The sample was trimmed with a 90° diamond knife (Diatome) to expose the cells at two surfaces (the imaging surface and the surface perpendicular to the focused ion beam, FIB) and optimize the acquisition (55). When targeting the symbiotic microalgae within its host, the trimming was targeted towards the periphery of the organisms where they were in higher numbers. After the sample was trimmed, it was mounted onto the edge of a SEM stub (Agar Scientific) using silver conductive epoxy (CircuitWorks) with the trimmed surfaces facing up and towards the edge of the stub. The sample was gold sputter coated (Quorum Q150RS; 180 s at 30 mA) and placed into the FIB-SEM for acquisition (Crossbeam 540, Carl Zeiss Microscopy GmbH). Once the ROI was located in the sample, Atlas3D software (Fibics Inc. and Carl Zeiss Microscopy GmbH) was used to perform sample preparation and 3D acquisitions. First, a 1 μm platinum protective coat (20-30 μm^2^ depending on ROI) was deposited with a 1.5 nA FIB current. The rough trench was then milled to expose the imaging cross-section with a 15 nA FIB current, followed by a polish at 7 nA. The 3D acquisition milling was done with a 1.5 nA FIB current. For SEM imaging, the beam was operated at 1.5 kV/700 pA in analytic mode using an EsB detector (1.1 kV collector voltage) at a dwell time of 8 μs with no line averaging. For each slice, a thickness of 8◻nm was removed, and the SEM images were recorded with a pixel size of 8◻nm, providing an isotropic voxel size of 8 × 8 × 8 nm^3^. Whole volumes were imaged with 1000–1200 frames, depending on the *Brandtodinium* cells. Raw FIB-SEM stacks are available at https://www.ebi.ac.uk/biostudies/studies/XXXXX. Source data are provided with this paper.

The first step of image processing was to crop the free-living and symbiotic *Brandtodinium* cells using the open software Fiji (https://imagej.net/Fiji), followed by image registration (stack alignment), noise reduction, semi-automatic segmentation, 3D reconstruction of microalgae cells and morphometric analysis as in (17). Image registration was done by the FIJI plugin ‘Linear Stack Alignment with SIFT’(56), then fine-tuned by AMST (57). Aligned image stacks were filtered to remove noise and highlight contours using a Mean filter in Fiji (0.5 pixel radius). Segmentation of organelles and other cellular compartments of *Brandtodinium* and Collodaria was carried out with 3D Slicer software (58) (https://www.slicer.org), using a manually-curated, semiautomatic pixel clustering mode (5 to 10 slices are segmented simultaneously in z) as in (17). We assigned colours to segmented regions using paint tools and adjusted the threshold range for image intensity values. Morphometric analyses were performed with the 3D slicer module “segmentStatistics” on the different segments (segmented organelles) and converted to μm^3^ or μm^2^ taking into account the voxel size of 8 nm (Table S1).

### Synchrotron X-ray fluorescence (S-XRF) imaging

S-XRF hyperspectral images were acquired on the ID21 and ID16B-NA beamlines of the European Synchrotron Radiation Facility (Cotte et al. 2017, Martinez-Criado et al. 2016). 300 nm-thick cell sections were laid on Si_3_N_4_ membranes. On ID21 the incoming X-rays were tuned to the energy of 7.3 keV with a fixed-exit double crystal Si (111) monochromator, and focused to 0.3×0.8 μm^2^ with a Kirkpatrick-Baez (KB) mirror system, yielding a flux of 5·10^10^ ph/s. The experiment was performed under vacuum (10^−5^-10^−4^ mbar). The emitted fluorescence signal was detected with energy-dispersive, large area (80 mm^2^) SDD detectors equipped with a Be window (SGX from RaySpec). Images were acquired by raster-scanning the sample in the X-ray focal plane, with a 0.5×0.5 μm^2^ step and 500 ms dwell time. The detector response was calibrated over a thin film reference sample consisting of layers of elements in ng/mm^2^ concentration sputtered on a 200 nm thick Si_3_N_4_ membrane (RF7-200-S2371 from AXO), measured using the same acquisition parameters.

On ID16B-NA, a beam of 17.5 keV focused to 50×50 nm^2^ through KB mirrors was used to excite the samples. The photon flux at sample was ~ 2·10^11^ ph/s. High-resolution XRF images (50×50 nm^2^ step size) were acquired in air, with a dwell time of 100 ms/pixel. Two 3-element SDD detector arrays were used to collect fluorescence from the sample. The detector response was calibrated over a thin film reference sample (RF8-200-S2453 from AXO). High-resolution images were acquired for free and symbiotic microalgae, and in selected areas of the hosts.

Hyperspectral images acquired in both beamlines were normalized by the incoming photon flux and subjected to the same data analysis protocol. Elemental mass fractions were calculated from fundamental parameters with the PyMca software package (60), assuming a biological matrix of light elements (H, C, N, O) and a density of 1 g/cm^3^, as reported in the NIST Star database for the standard composition of soft tissue (https://www.physics.nist.gov/cgi-bin/Star/compos.pl?matno=261). The calculation of the elemental concentrations in specific areas were performed by manually selecting the pixels in the region of interest and summing up their fluorescence signal; the sum spectrum normalized by the number of pixels was then subjected to spectral deconvolution, and the peak areas were converted in mass fractions (See also Table S4).

### NanoSIMS mapping of C, N, P and S in samples of natural isotopic composition

The wafers containing thin sections were first coated with 20 nm gold-palladium and analyzed with a NanoSIMS 50L (Cameca, Gennevilliers, France), using 2 pA beam of 16 keV Cs^+^ primary ions focused to 70 nm spot. The analyzed sample area of 25×25 μm² was scanned in 512 × 512 pixel raster with a dwelling time of 2 ms/pixel. Prior the analysis, the area of 100×100 μm² involving the analysis FoV was pre-implanted for 15 min with 200 pA Cs^+^ beam to equilibrate the secondary ion yield. The data were acquired in 20-30 plains upon consecutive scanning with the primary ion beam. Secondary ions extracted from each pixel of the sample surface (^16^O^−^, ^12^C_2_^−^, ^12^C^13^C^−^, ^12^C^14^N^−^, ^13^C^14^N^−^, ^31^P^−^ and ^32^S^−^) were separated according to their mass to charge ratio (*m/z*) with mass resolving power above 8000 (MRP=M/dR) achieved with 20×140 μm nominal size (width×height) of the entrance slit, 200×200 μm aperture, 40×1800 μm exit slits and the energy slit blocking 20% of ions at their high-energy distribution site. Complete 200 nm of section depth was consumed upon the analysis. The acquired data were evaluated using Look@nanoSIMS (LANS) software (61).

### ^13^C- and ^15^N-labeling for TEM-NanoSIMS analysis

Collodarians were collected at the subsurface in the Mediterranean Sea (Villefranche-sur-Mer, France) in September 2019. Live collodarians were then isolated under a binocular, transferred in 50 mL plastic flasks filled with filtered (0.22 μm) natural seawater (collected at the sea surface the same day). The day after the sampling, the incubation was initiated at 2pm under controlled light and temperature conditions (20°C) by spiking with H^13^CO_3_^−^ and ^15^NH_4_^+^ (99%, Sigma− Aldrich) to reach final concentrations of 2 mM and 10 μM, respectively. After 3h of incubation, the experiment was stopped by removing the collodarian colonies from the spiked seawater and transferring them in filtrated seawater. The samples were then immediately cryofixed with high pressure freezing and processed as explained above. For correlated EM - NanoSIMS, ultrathin sections of 70 or 200 nm thick were obtained. The 70 nm sections were mounted onto copper grids or slots coated with a formvar - carbon film. Sections were then stained in 1% uranyl acetate (10min) and lead citrate (5min). Micrographs were obtained using a Tecnai G2 Spirit BioTwin microscope (FEI) operating at 120 kV with an Orius SC1000 CCD camera (Gatan). The 200 nm sections were mounted on silicon wafers and imaged with a GeminiSEM (Zeiss) operating in ESB mode at 3 kV. For NanoSIMS observations, sections were coated with 10 nm of gold and analyzed with a NanoSIMS 50L (Cameca, Lausanne, Switzerland), using a cesium (Cs^+^) primary ion beam with a current of about 2 pA and an energy of 16 keV. The primary ion beam was focused to a nominal spot size of ~ 120-150 nm and stepped over the sample in a 256×256 pixel raster to generate secondary ions with a dwelling time of 5 ms/pixel. The raster areas (scanning surface area) were defined based on previous SEM/TEM observations and were between 10×10 μm^2^ to 40×40 μm^2^. Secondary ions (^19^F^−^, ^12^C_2_^−^, ^13^C^12^C^−^, ^12^C^14^N^−^, ^12^C^15^N^−^, ^31^P^−^ and ^32^S^−^) were simultaneously collected in electron multipliers at a mass resolution (M/ΔM) of about 8000 (Cameca definition). Each NanoSIMS image consist of 6 to 10 sequential images, drift corrected, and accumulated using the software Look@NanoSIMS (61) to quantify mean ^13^C and ^15^N enrichments of different sub-cellular structures. For control (i.e. not enriched) samples, four images were acquired including 5 symbiotic dinoflagellates and two different areas of the host cell (Table S2). These control images were used to determine the standard ratio against which the experimental ^13^C-and ^15^N-enrichments were quantified. For ^13^C and ^15^N-enriched samples, two distinct collodarian colonies were analyzed.

### Quantitation of assimilation activity

The cellular activity in assimilation of carbon and nitrogen was derived from the detected changes in nitrogen and carbon isotopic composition. Changes in ^13^C and ^15^N fractions are not directly proportional to cellular activity and the assimilation activity was therefore expressed as the relative assimilation K^i^_A_ (29):

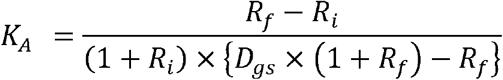

The K^i^_A_ represents the *fraction* (in at%) of carbon (^12^C and ^13^C) or nitrogen (^14^N and ^15^N) assimilated from a growth substrate into a biomass unit *relatively to its initial* content (carbon or nitrogen; before incubation with a labelled substrate) in a biomass unit e.g. cell or cell fragment.

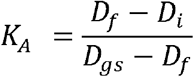

*R*_*i*_ / *D*_*i*_ – initial isotope ratio/fraction in biomass unit (before incubation with isotope-labelled growth substrate)

*R*_*f*_ / *D*_*f*_ – final isotope ratio/fraction in biomass unit (after incubation with isotope-labelled growth substrate)

*D*_*gs*_ – fraction of heavy isotope in growth substrate

### Bulk quantification of metals in collodaria (metallome)

Free-living microalgae (*Brandtodinium*) in culture and collodarian colonies (single colonies and a mix of different colonies) collected in surface waters of the Mediterranean Sea were centrifugated to remove the medium and seawater, respectively. Cells were then cryofixed and preserved at −80°C. For metal extraction, samples were dehydrated and mineralized in 200 μL 65% (w/v) ultrapure HNO_3_ at 90°C for 4 hours. Digested samples were diluted in distilled water and analyzed using an iCAP RQ quadrupole mass instrument (Thermo Fisher Scientific GmbH, Germany). The instrument was used with a MicroMist U-Series glass concentric nebulizer, a quartz spray chamber cooled at 3°C, a Qnova quartz torch, a nickel sample cone, and a nickel skimmer cone equipped with a high-sensitivity insert. Elements were analyzed using either the standard mode (for ^27^Al, ^31^P, ^111^Cd) and/or the kinetic energy discrimination mode with helium as the collision cell gas (for ^31^P, ^55^Mn, ^56^Fe, ^57^Fe, ^58^Ni, ^59^Co, ^60^Ni, ^63^Cu, ^64^Zn, ^65^Cu, ^66^Zn). Concentrations were determined using standard curves and corrected using an internal standard solution of 103Rh added online. Data integration was done using the Qtegra software (version 2.8.2944.115). Element content was normalized to phosphorus (P) content as in (45). (See Table S5)

