## supplementary figures for "Subcellular architecture and metabolic connection in the planktonic photosymbiosis between Collodaria (radiolarians) and their microalgae"

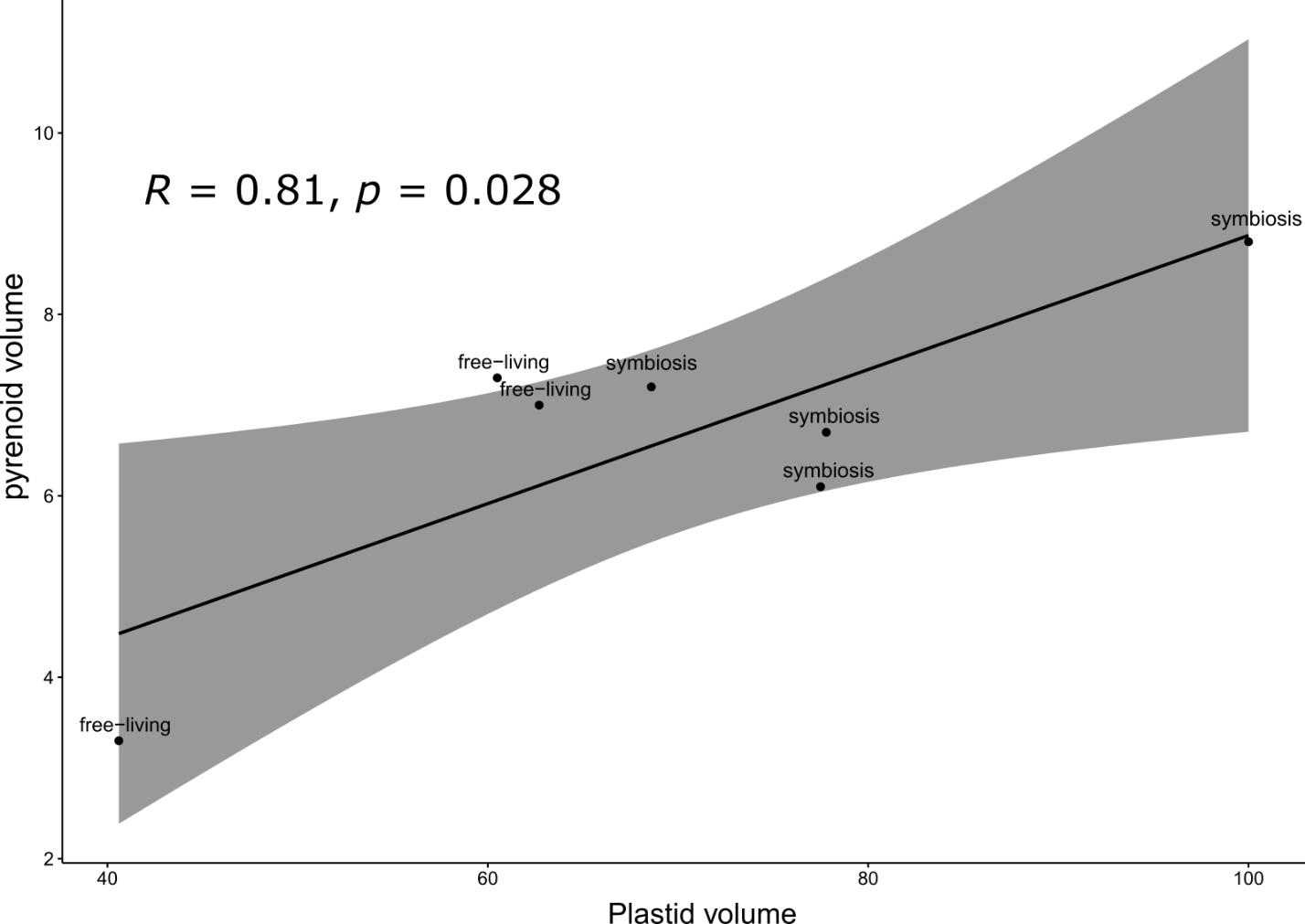

**Figure S1:** Correlation between volume of pyrenoid and volume of plastids (µm^3^) of free-living and symbiotic microalgae (*Brandtodinium*) after FIB-SEM imaging and 3D reconstruction.

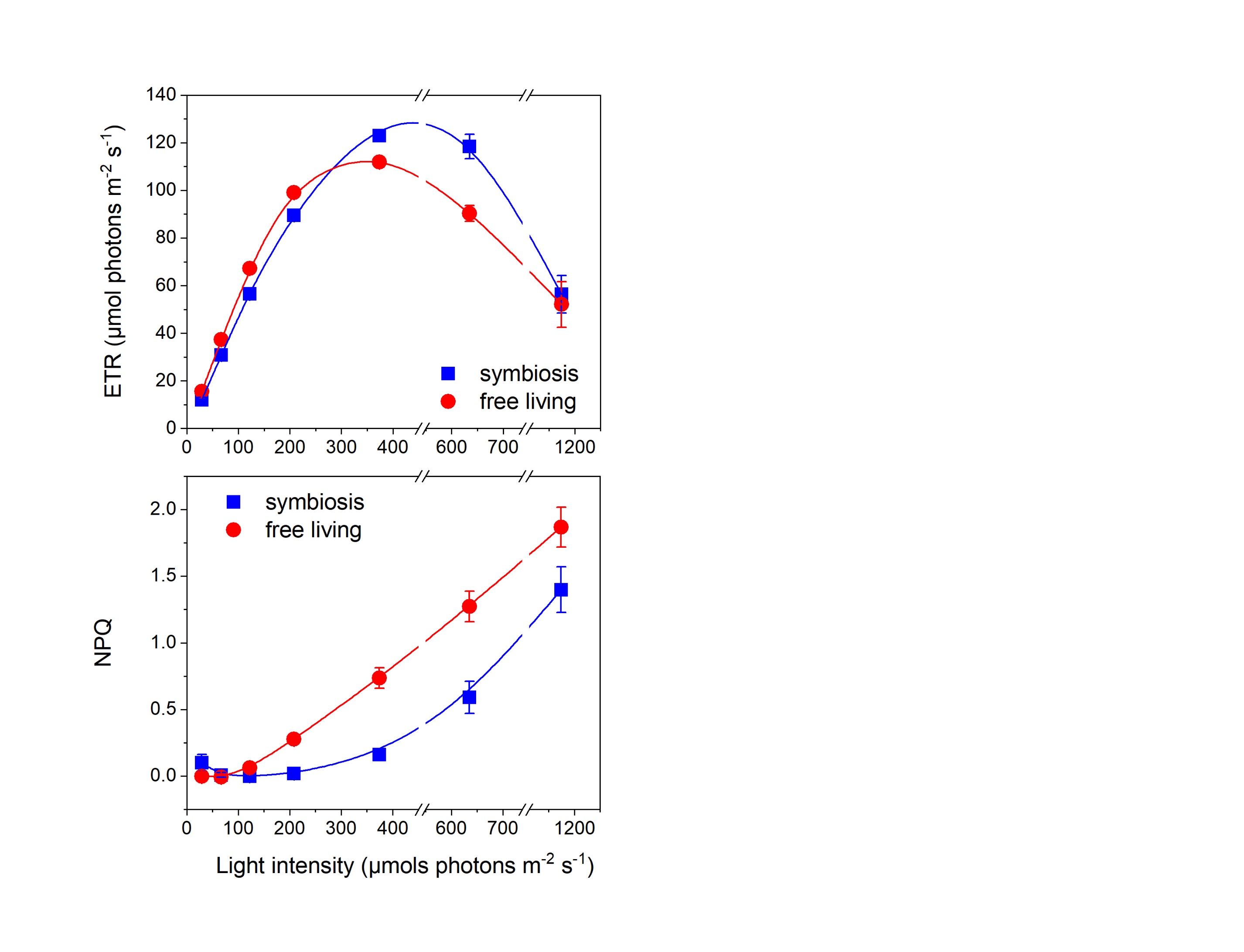

**Figure S2:** The non-photochemical quenching (NPQ) parameter indicated that the light energy absorption of symbiotic microalgae *Brandtodinium* (blue line; n=4) was less sensitive in high-light conditions than it was in free-living phase grown in culture (red line; n=4).

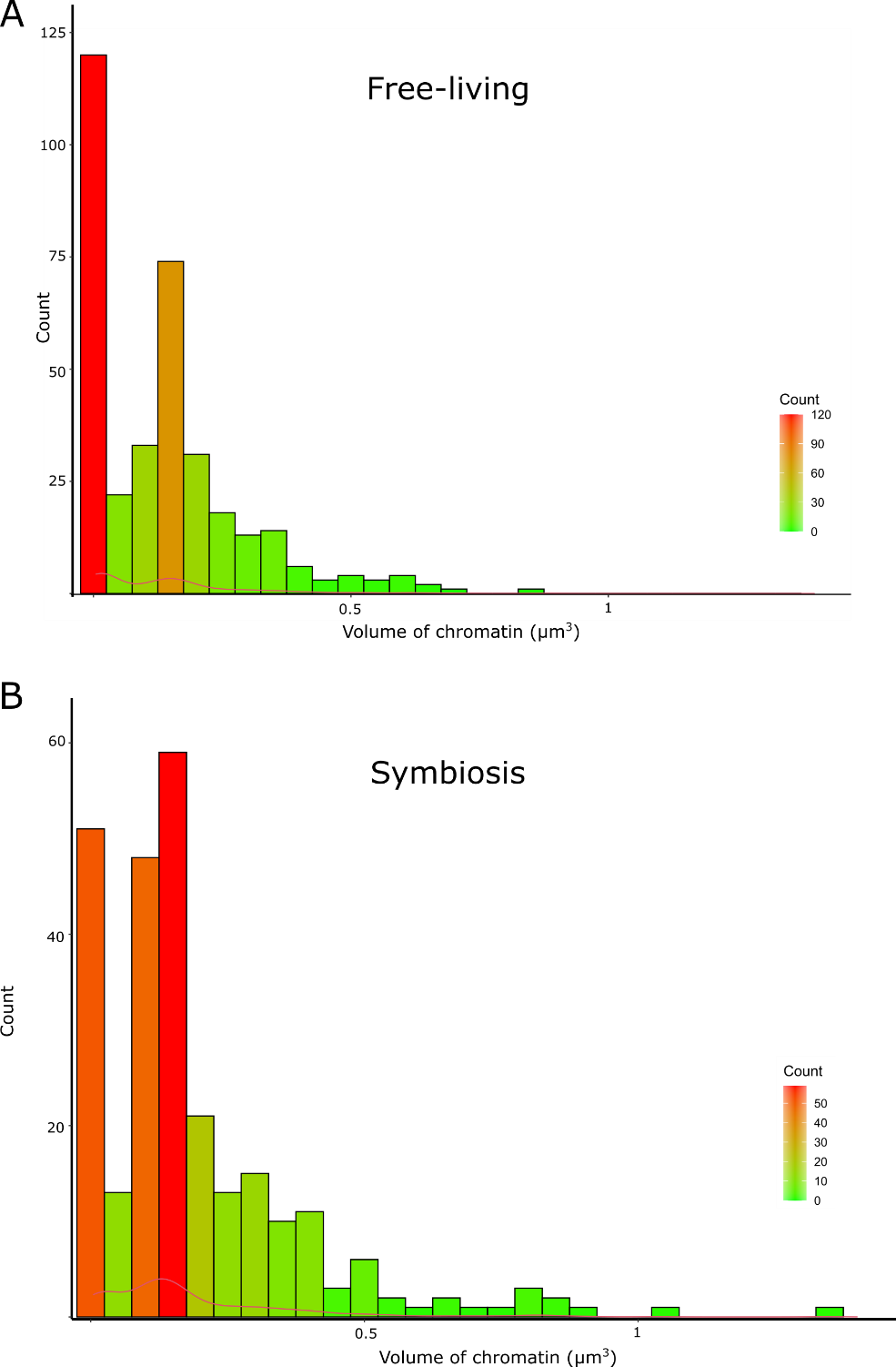

**Figure S3:** Distribution of heterochromatin volume of free-living and symbiotic microalgae *Brandtodinium* (color coded: red bars correspond to high numbers of heterochromatin). Heterochromatin, which includes rod-shaped chromosomes and condensed chromatin present in the nucleus, has been reconstructed in 3D after FIB-SEM imaging and its volume quantified. In free-living, we observed a high number of protruding small condensed chromatin structures in the nucleus compared to symbiosis.

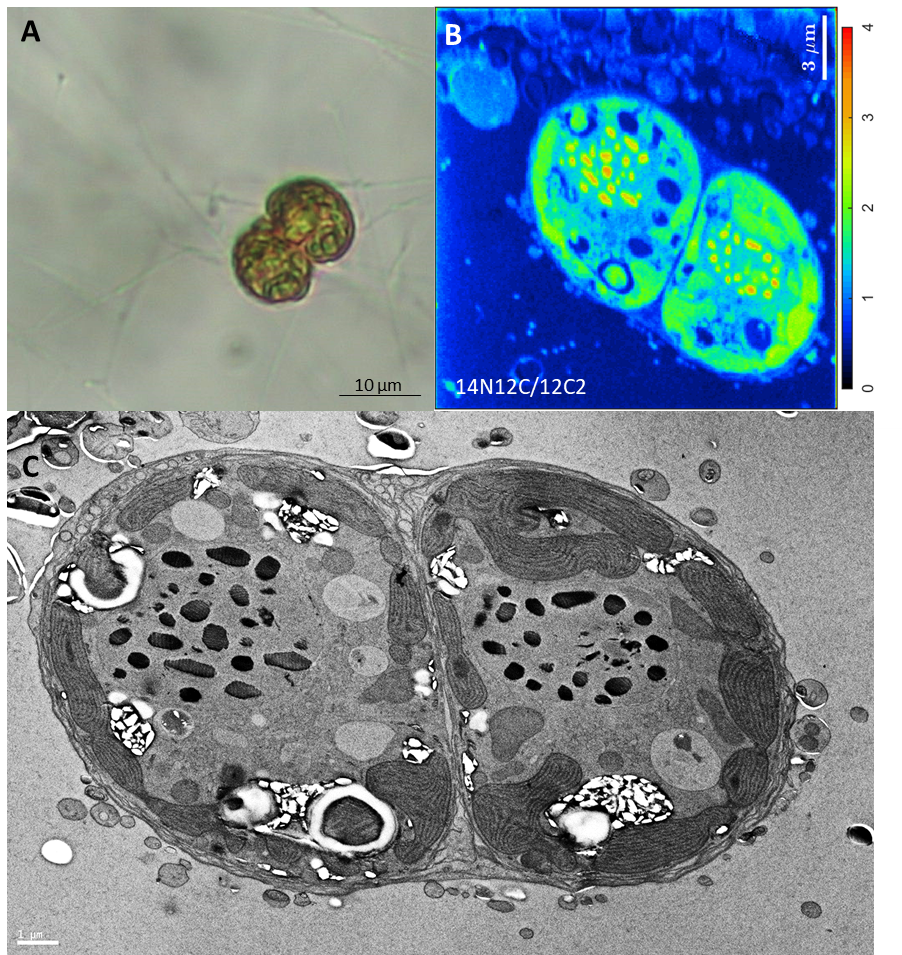

**Figure S4:** Observation with light microscopy (A), nanoSIMS (^14^N^12^C/^12^C_2_ ratio) (B) and Transmission Electron microscopy (C) of dividing microalgal cells (*Brandtodinium*) embedded in the gelatinous matrix of the Collodaria (host).

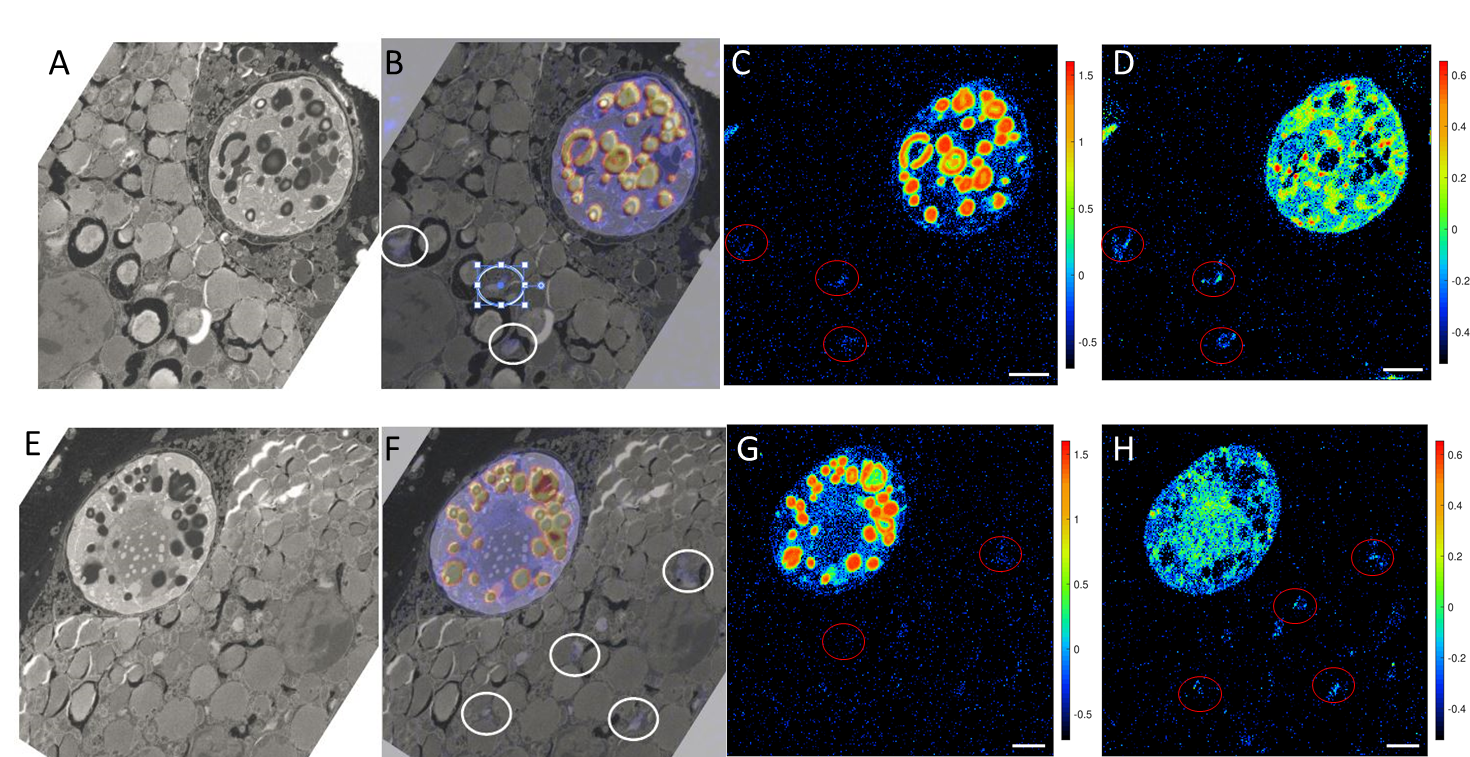

**Figure S5:** Correlated Transmission Electron microscopy (A and B) with nanoSIMS images showing relative assimilation (log(ka)) of carbon (C and G) and nitrogen (D and H) in symbiotic microalgae *Brandtodinium* and in the Golgi apparatus (highlighted by a red circle) of the host cell (Collodaria).

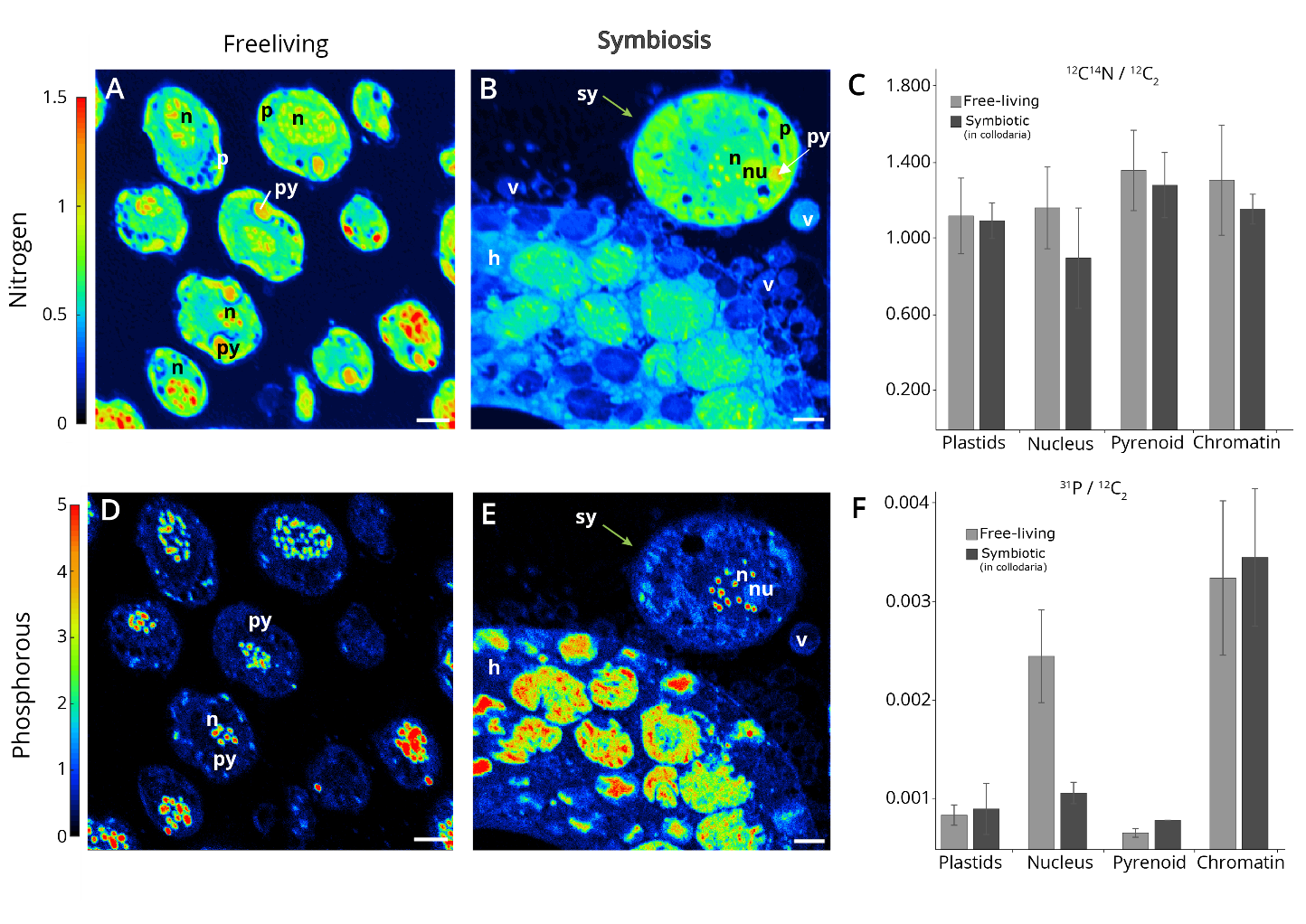

**Figure S6:** Subcellular quantitative mapping with nanoSIMS of natural nitrogen and phosphorous in the free-living and symbiotic stage of *Brandtodinium*. **A and B:** Nitrogen (^12^C^14^N/^12^C_2_) mapping in the free-living (A) and symbiotic (B) stages. Note that part the host cell is visible in B and the symbiotic microalga is indicated by a green arrow. **D and E**: Phosphorous (^31^P/^12^C_2_) mapping in the free-living (D) and symbiotic (E, green arrow) stages. Nitrogen (^12^C^14^N/^12^C_2_) and phosphorous (^31^P/^12^C_2_) content in different organelles (plastids, nucleus, pyrenoid and chromatin) of free-living (light green) and symbiotic (dark green) microalgae. (n: nucleus; nu: nucleolus; py: pyrenoid; p: plastid; h: host cell (Collodaria); m: matrix; v: vacuoles in the matrix) (See also Table S3).

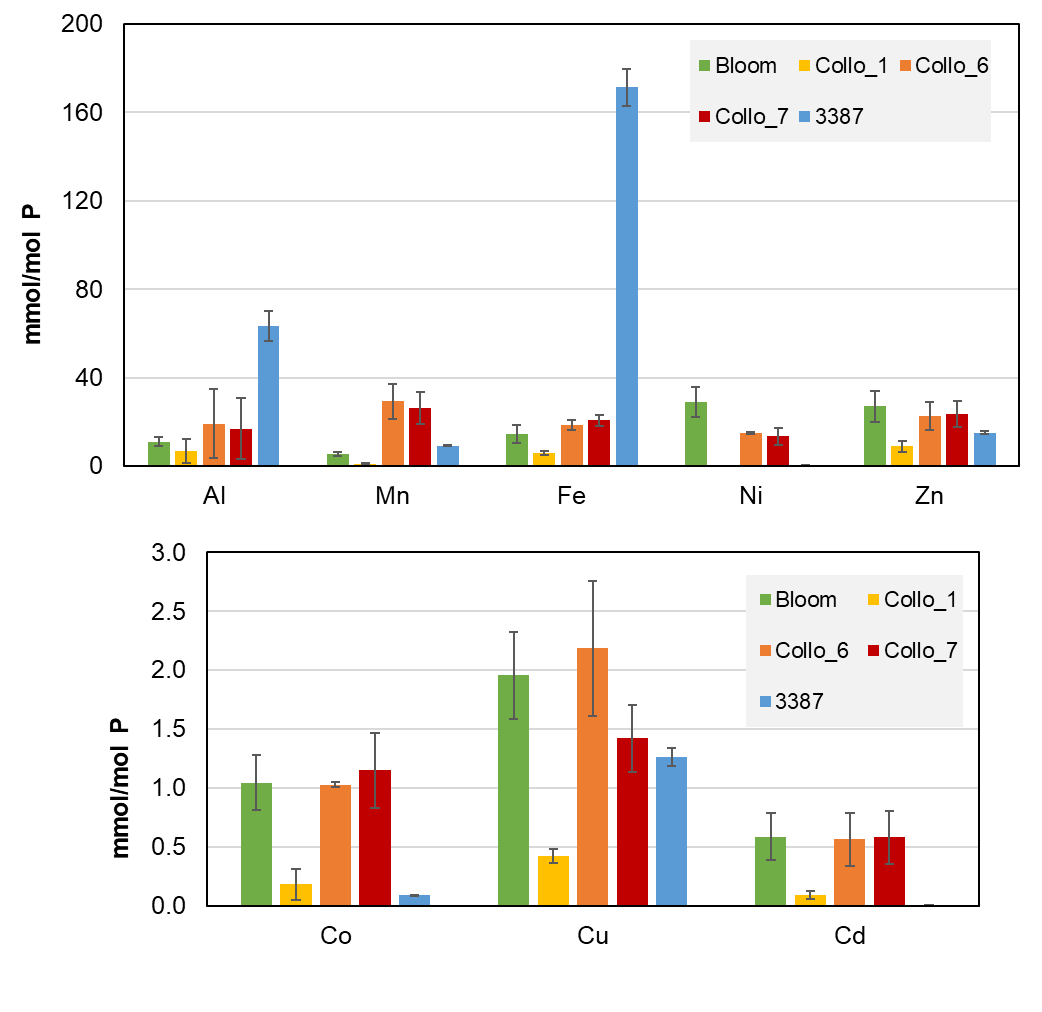

**Figure S7:** Metallome analyses of Collodaria (containing symbiotic microalgae, *Brandtodinium*) showing the intracellular metal quotas normalized against phosphorus as the biomass indicator (mmol of metal/mol of P). Al: Aluminum, Mn: Manganese, Fe: iron, Ni: nickel, Zn: Z=zinc, Co: cobalt, Cu: copper, Cd: cadmium. Bloom sample correspond to a pool of different collodarian colonies, Collo 1, 6 and 7 are individual colonies and sample 3387 represents the free-living microalgae *Brandtodinium* from culture medium. (See also Table S5).

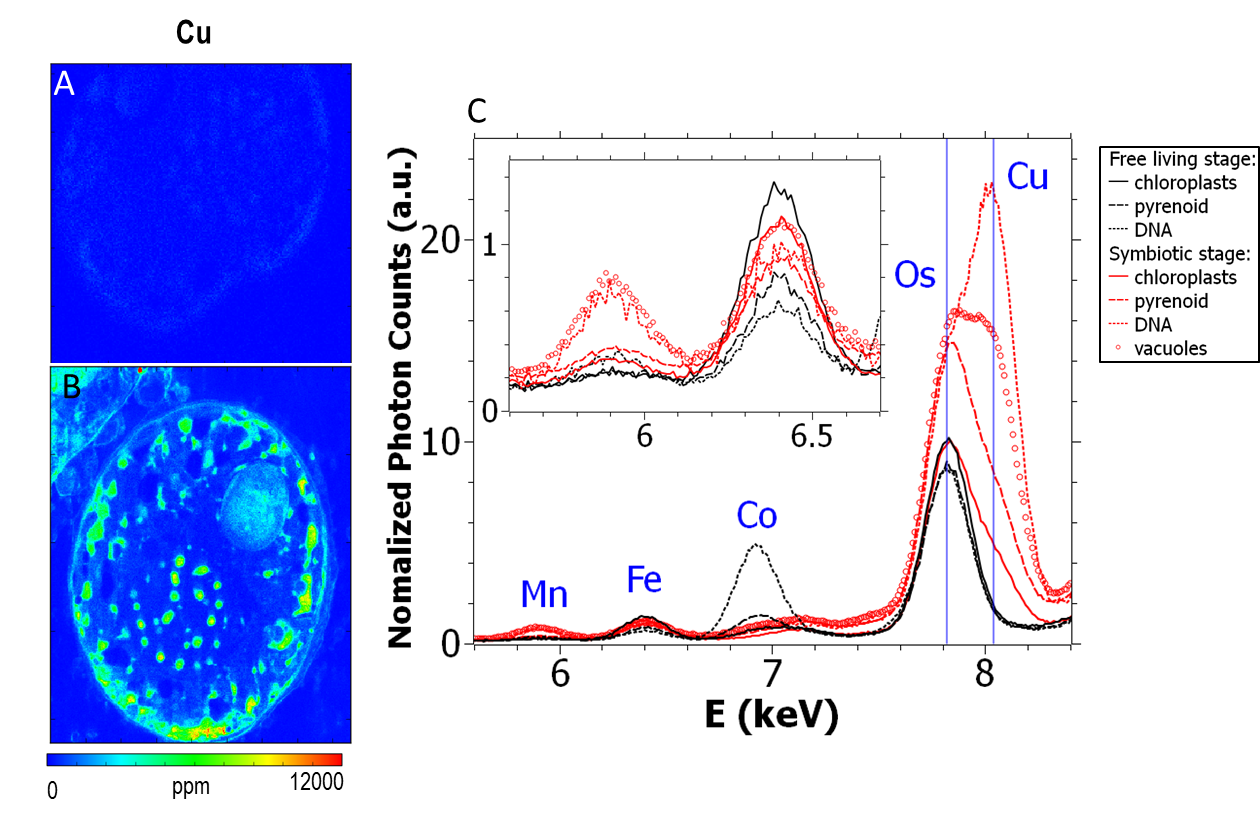

**Figure S8:** Subcellular quantitative mapping using S-XRF (Synchrotron X-ray Fluorescence) of copper (Cu) showing its concentration (ppm) in free-living (**A**), and symbiotic microalga *Brandtodinium* (**B**) with a xy resolution of 50 nm and at 7.3 keV. Note that Cu was below the detection level in free-living while in symbiosis it was concentrated in condensed chromatin and vacuoles closed to plastids. **C:** Average S-XRF spectrum per pixel in subcellular compartments (chloroplasts, pyrenoid and condensed chromatin or DNA) of *Brandtodinium* in free-living (black lines) and symbiotic (red lines) stages, showing high Cu concentration in condensed chromatin and vacuoles of symbiotic microalgae. Osmium peak is not interfering with Cu as such concentration.
